# Glutamine Tautomerization Drives RhoGAP‑Aided GTP Hydrolysis in Small Rho GTPases

**DOI:** 10.64898/2026.03.18.712581

**Authors:** Angela Parise, Riccardo Rozza, Karolina Mitusinska, Alessandra Magistrato

## Abstract

Rho GTPases promote GTP hydrolysis aided by specific GTPase-activating proteins (GAPs). By alternating between an active GTP-bound and an inactive GDP-bound state, Rho GTPases function as molecular switches regulating cytoskeletal dynamics and cell motility. Despite their biological relevance, the detailed molecular mechanism underlying Rho GTPases catalysis remains contentious. Here, using classical and hybrid quantum–classical molecular dynamics, we resolve the mechanism of GTP hydrolysis in the RhoGAP-RhoA complex. We reveal that GTP hydrolysis proceeds through a dissociative nucleophilic substitution mechanism, driven by an amide → imide tautomerization of Gln63, which aids in delivering a proton from the nucleophilic water to the leaving phosphate group. The Gln63 imide tautomer also loosens RhoGAP–RhoA interfacial contacts, allowing solvent molecules to enter and drive a water-mediated reverse tautomerization of Gln63 that restores the catalytically-competent configuration of the RhoA active site. Conservation of key interface residues across Rho/Rho GAP family members suggests that this mechanism may be shared by most Rho GTPases.

## Introduction

Human small GTP-binding proteins of the Ras superfamily coordinate diverse cellular functions, including cytoskeletal reorganization, vesicular trafficking, cell proliferation, and motility.^1–3^ Specifically, members of the Rho family govern cytoskeletal rearrangements: Rac1 (Ras-related C3 botulinum toxin substrate 1) promotes lamellipodia formation via actin polymerization, Cdc42 (cell division control protein 42 homolog) drives filopodia extension, and RhoA (Ras homolog family member A) stimulates stress fiber and focal adhesion assembly.^4^ Viruses hijack these pathways to enhance infectivity,^5^ while abnormal activation or misregulation of Rho GTPases function is implicated in tumor progression and metastasis.^4,6–8^ Elucidating the molecular mechanisms of Rho GTPases is therefore biologically significant and offers tangible opportunities for therapeutic development. ^9^

Small Rho GTPases function as molecular switches, cycling between a GTP-bound (active) and a GDP-bound (inactive) state, ^10^ regulated by the action of GTPase-activating proteins (GAPs), which promote GTP hydrolysis, and by guanine nucleotide exchange factors (GEFs) proteins, which facilitate GDP release. ^11^ At structural level, Rho GTPases exhibits two functional regions within their G-domain. These are the so-called switch I and switch II (Figure 1A), which shape the substrate-binding pocket, drive nucleotide-dependent conformational changes, and mediate interactions with GTPase regulators, including GEF and GAP proteins.^12–14^

**Figure 1.**
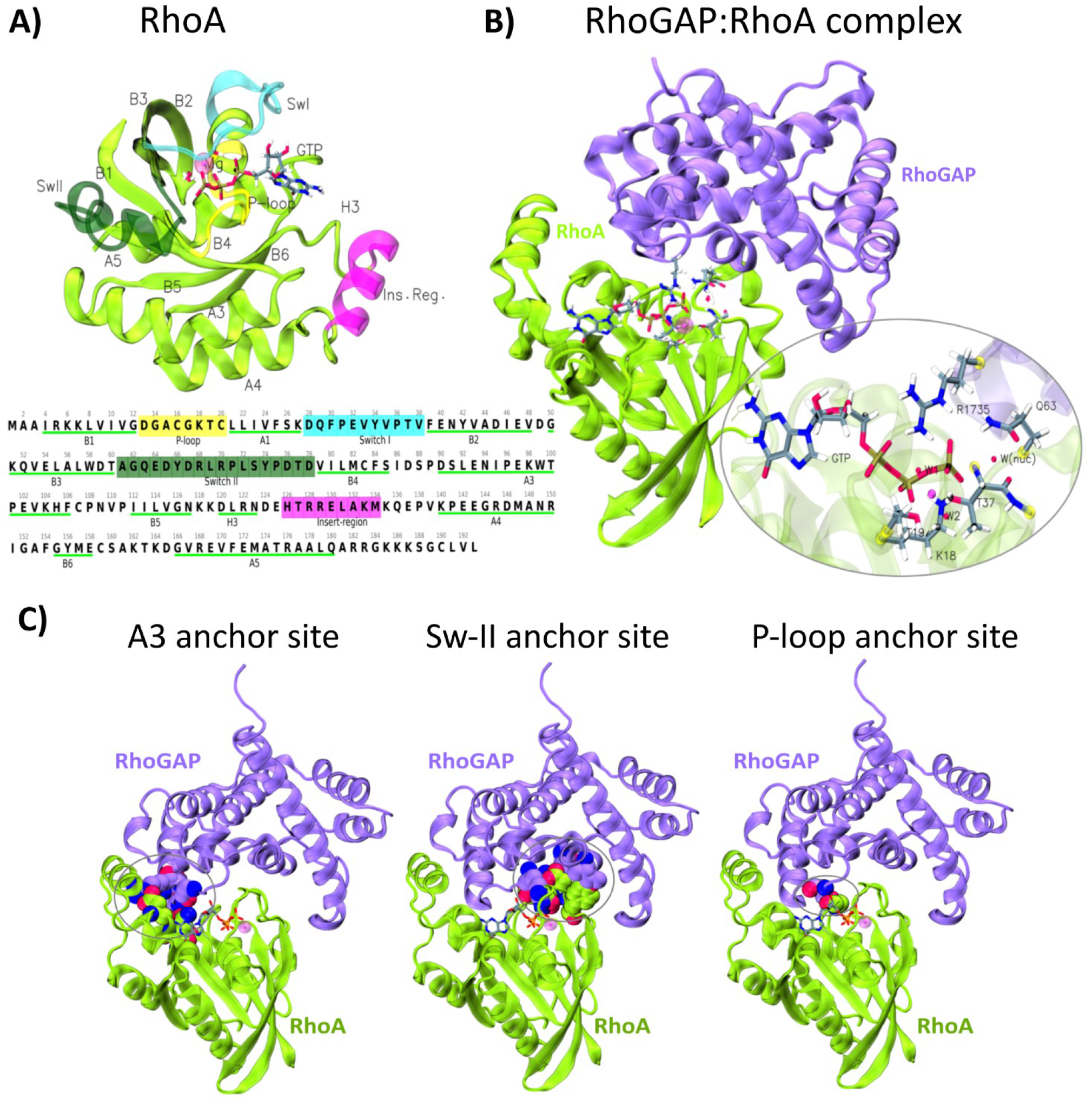
Structure of GTP-bound RhoA and RhoGAP:RhoA complex. A) Cartoon representation (top) and primary sequence (bottom) of RhoA. The main functional motifs: switch I, switch II, p-loop, and insert region, are highlighted in cyan, dark green, yellow, and purple, respectively. The Mg^2+^ ion is shown as a magenta sphere, and the substrate (GTP) is shown in sticks. B) Cartoon representation of GTP-bound RhoA (green) in complex with the RhoGAP domain of human Myosin 9b (Myo9b-RhoGAP, purple). The active site is circled, key interface residues involved in GTP hydrolysis are shown in sticks. For clarity, hydrogen atoms of water molecules are omitted. C) Key interactions at the interface. The three main anchoring sites are: (i) A3 site, where RhoGAP residues Gly1738, Ala1740, Arg1742, and Arg1744 interact with RhoA helix A3 residues Asn94, Asp90, and Asp97; (ii) switch II site, where RhoGAP residues Arg1735 (arginine finger), Arg1776, and Ile1848 interact with RhoA switch II residues Gln63 (catalytic), Glu64, Asp65, and Tyr66; (iii) P-loop site, involving RhoGAP residue Ser1737 interacting with Gly14 in the RhoA P-loop region. Only heavy atoms are shown in the van der Waals representation. Carbon atoms are colored in purple, green and gray for RhoGAP, RhoA residues, and GDP, respectively. Oxygen and nitrogen atoms are colored red and blue, respectively.

Spontaneous GTP hydrolysis proceeds extremely slowly in water^15^ and remains inefficient even when catalysed by isolated Rho GTPases (i.e. RhoA GTP hydrolysis rate is about 0.022 min^−1^ at 300 K,^16^ which corresponds to an activation free energy barrier (ΔG^‡^) of about 23 kcal·mol^−1^ according to Eyring equation^17^ within the framework of transition-state theory (TST)^18^). GAP proteins enhance Rho GTPases catalysis, significantly accelerating GTP hydrolysis.^19–21^ As an example, the specific GAP domain of RhoA (RhoGAP domain) accelerates GTP hydrolysis rate to ∼60 min^−1, 16^ corresponding to a ΔG^‡^ of ∼17.2 kcal·mol^−1^, by inserting a conserved arginine—known as the “arginine finger”—into the GTPase active site. ^22^ The arginine finger optimally orients the catalytic water and the key Gln residue, conserved in Rho GTPases, while also stabilizing the negative charge that accumulates on the γ-phosphate at the transition state (Figure 1B).^23,24^

Despite many high-resolution structural data and extensive biochemical and computational studies,^9,25–28^ the catalytic mechanism of Rho GTPases is still a matter of debate. Different mechanism have been proposed (Scheme S1): *(i)* the *solvent-assisted pathway* in which the deprotonation of the nucleophilic water occurs after the nucleophilic attack, without support of a general base; *(ii)* the *substrate-assisted pathway*, where the γ-phosphate of GTP accepts the proton released by the nucleophilic water as it performs the attack; *(iii) the general base pathway*, in which proton abstraction from the nucleophilic water is supported by a conserved glutamine residue (e.g., Gln63 in RhoA) or by an additional water molecule (two waters mechanism); *(iv)* the *tautomerism-driven pathway*, where an amide → imide tautomerization of the Glutamine side chain occurs. ^21^

Numerous computational studies have explored these mechanisms across multiple small GTPases, ^29–35^ but only a few obtained results consistent with experimentally measured kinetic constants. Moreover, in some studies restoration of the catalytic site was unexpectedly identified as the rate-limiting step. ^32,35^

To uncover the molecular mechanism of GTP hydrolysis in RhoA and to determine whether it extends to other Rho GTPases, we used an integrated approach based on classical (MM) and hybrid quantum– classical (QM/MM) molecular dynamics (MD) simulations. We reveal that GTP hydrolysis occurs via a dissociative nucleophilic substitution (S_N_1) reaction (Scheme S2). The rate-limiting step is the nucleophilic attack of the water molecule (W_nuc_) on the γ-phosphate, a process strongly facilitated by Gln63 through a tautomerization-mediated proton-shuttling process. We further identify a previously unrecognized route for restoring the active site into a catalytically competent state. Formation of the Gln63 imide tautomer weakens RhoGAP-RhoA interfacial contacts, and allowing solvent molecules to access the catalytic site, and thus promoting water-assisted imide→ amide reverse tautomerization of Gln63 via a low-energy barrier route. Bioinformatic and structural analysis of diverse Rho GTPase–GAP complexes reveal broad conservation of both catalytic residues and those implicated in Gln-imide–induced decoupling of RhoGAP–RhoA indicates that this tautomerization-based mechanism is likely operative throughout most Rho GTPase members.

## Methods

### System preparation

The initial coordinates of the system were obtained from the crystallographic structure of the RhoGAP:RhoA complex (PDB ID: 5HPY), which includes the full G-domain of RhoA in complex with the RhoGAP domain of human Myosin 9b, the catalytic Mg^2+^ ion, and a GTP analogue (GDP–F_3_Mg^−^). The native GTP molecule was reconstructed by replacing the corresponding atoms of the crystallized analogue using Maestro program (Schrödinger, LLC ^36^). Crystallographic water molecules, including the catalytic water (W_nuc_), were retained. Missing heavy atoms and crystallographically unresolved hydrogen atoms were added in sterically favorable positions. Protonation states of ionizable residues were assigned based on pK_a_ predictions from the PropKa 3 server, ^37,38^ with a reference pH of 7.5, consistent with experimental conditions used in kinetic assays.^16,19^ GTP was modelled as fully deprotonated according to IR and NMR data showing that GTP is known to be fully deprotonated in Ras^39^

### Classical MD Simulations

Topologies of RhoGAP:RhoA-GTP (reactant), RhoGAP:RhoA(Gln63*)-GDP+Pi (intermediate), and RhoGAP:RhoA-GDP+Pi (product) complexes were generated with the LEaP module of AmberTools23 ^40^. The protein was described with the ff14SB force field^41^ and solvated in a rectangular box of TIP3P water molecules, ^42^ with a 14 Å buffer between the solute and the box edges. Parameters for GDP and GTP were derived by combining AMBER94 force field data for guanosine monophosphate with parameters for the terminal phosphate groups of adenosine di- and triphosphate from the AMBER parameter database. To neutralize the system, 11 Na^+^ counterions were added, described by the Joung & Cheatham parameters, ^43^ optimized for monovalent ions in TIP3P water. The catalytic Mg^2+^ ion was modeled using parameters from Li *et al*. ^44^. Despite known limitations of classical force fields for magnesium ions, due to lack of polarization and charge-transfer effects in the presence of negatively charged phosphates, ^45^ these parameters are suitable for comparative studies as shown in our previous investigations. ^8,13,14^ The parameters of the imide form of glutamine residue (Gln63*), were generated by cleaving the Cα–Cβ bond and adding a hydrogen atom (H_L_) to complete the Cβ valence. Geometry optimizations were performed at the DFT-B3LYP/6-311G* level using Gaussian16. ^46^ Partial atomic charges were derived using the RESP method by fitting HF/6-31G* electrostatic potentials with the Merz–Singh–Kollman scheme. The parameters of the inorganic phosphate (Pi) were derived consistently using the same DFT optimization and RESP protocol. AMBER force field parameters for both Gln63* and Pi were assigned via Antechamber (AmberTools23 ^40^). The backbone atom charges of Gln63* were restrained to ff14SB values to ensure compatibility with the protein environment. Overall, four system topologies were generated to represent the full catalytic cycle: (i) the reactant state (RhoGAP:RhoA-GTP, REA), (ii) the intermediate (RhoGAP:RhoA(Gln63*)-GDP+Pi) in its closed conformation (INT3), (iii) the intermediate in its open conformation (INT4), and (iv) the product state (RhoGAP:RhoA-GDP+Pi, PROD). Except for the reactant state, which was derived from the crystallographic structure, all other states were taken from representative frames of the intermediates sampled during QM/MM metadynamics trajectories (Figure 2). The metadynamics protocol, and the criteria adopted for the identification of free-energy basins and the extraction of representative structures, are detailed below.

**Figure 2.**
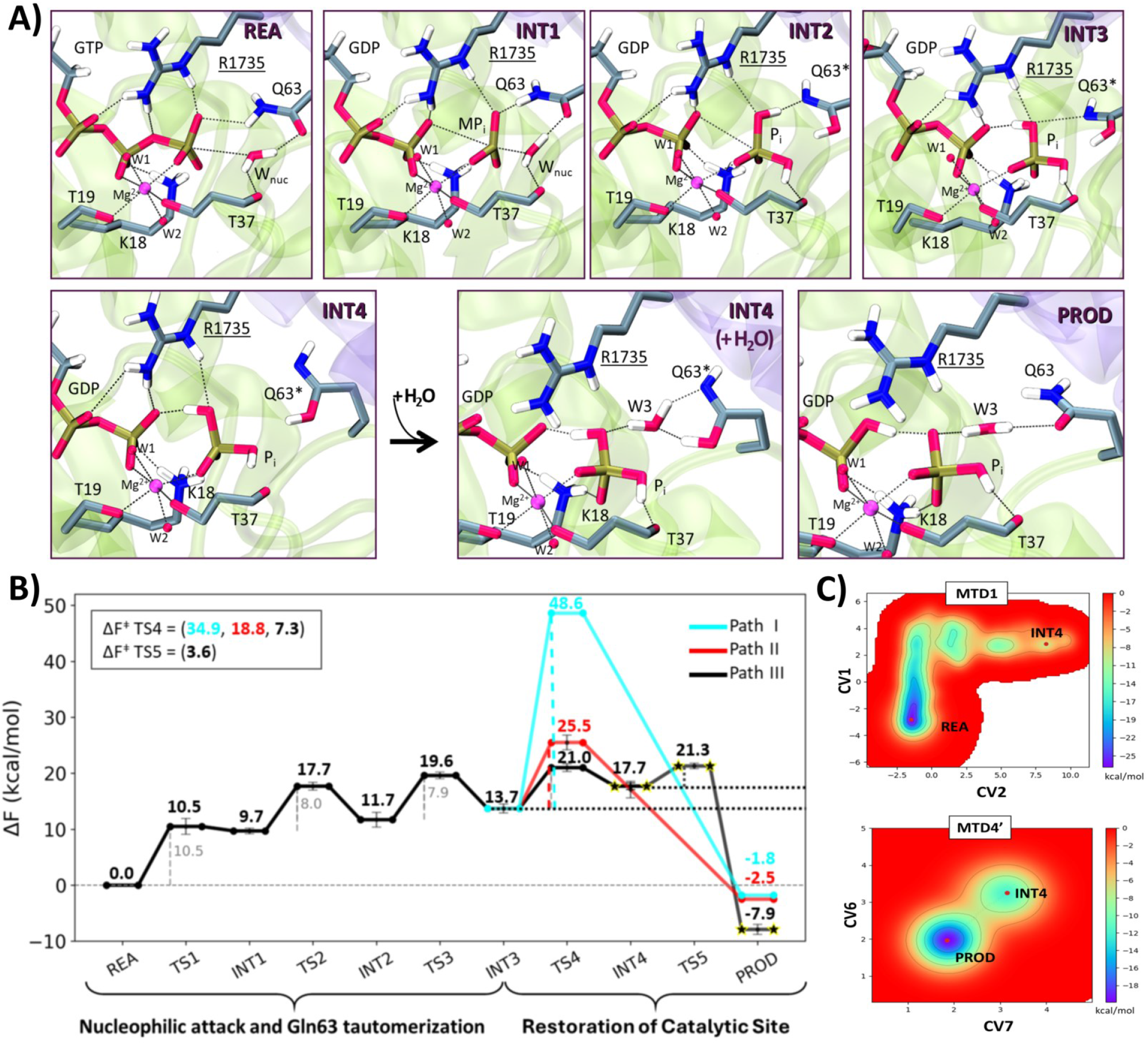
A) Representative structures of reactant, intermediate and product states of the GTP hydrolysis pathway in RhoGAP:RhoA complex. RhoA and RhoGAP are shown as transparent green and purple cartoons, respectively. Catalytic residues are shown as sticks and coloured by atom type. B) Schematic free-energy profile of GTP hydrolysis from QM(BLYP-D3(DZVP))/MM metadynamics simulations. The reaction proceeds in two phases: (i) nucleophilic attack and Gln63 tautomerization, leading to GDP and inorganic phosphate (Pi, INT2), followed by H-bond reorganization (INT3). Helmholtz free energies (ΔF) of minima and barriers up to INT3 are reported relative to the reactant state (kcal·mol^−1^). Restoration of the catalytic site, involving reverse tautomerization of Gln63* can occur via three pathways: intra-Gln63 proton transfer (Path I, cyan), phosphate-mediated restoration (Path II, red), and water-assisted restoration (Path III, stars). Error bars indicate the standard deviation over independent simulations. The three alternative pathways of Gln63* restoration are illustrated in Scheme S3, while the free-energy profiles of each pathway are reported in Figures S2– S5. C) Free-energy surface (FES, kcal·mol^−1^) of reaction Path III projected along the selected collective variables (CVs, atomic units). The FES highlights the starting and final states of MTD1 (top) and MTD4′ (bottom).

Each solvated structure was first energy-minimized with harmonic restraints (50 kcal·mol^−1^·Å^−2^) on all enzyme atoms using 5,000 steps of steepest descent followed by 5,000 steps of conjugate gradient. Hydrogen atom restraints were then released, and subsequent minimizations were performed with and without restraints on the protein backbone. Systems were gradually heated from 0 to 300 K over 3 ns in the NVT ensemble using the Langevin thermostat ^47^ with a 2 fs time step, followed by 2 ns equilibration in the NPT ensemble at 1 bar with the Berendsen barostat.^48^ For all MD simulations, the SHAKE algorithm was used to constrain light–heavy atom vibrations.^49,50^ Long-range electrostatics were treated with the particle mesh Ewald method ^51^ using a 12 Å cutoff, and harmonic restraints (20 kcal·mol^−1^·Å^−2^) were applied to maintain distances between Mg^2+^ and its coordinating water molecules. Production MD simulations of 0.5 μs were performed for the reactant, intermediates, and product states, with two additional independent 0.5 μs replicas for each system to assess reproducibility and statistical significance. All the simulations were run with the AMBER GPU program.^52^ Overall, we performed 6 µs of MD simulations.

### QM/MM MD

Unbiased QM/MM MD simulations were carried out for all minima along the reaction pathway using the CP2K program ^53^ (v. 2024.1). The MM region was described using the same force-field as in classical MD simulations, while the QM region was treated at the gradient-corrected Becke–Lee–Yang–Parr (BLYP) DFT level ^54,55^, supplemented with D3 dispersion corrections ^56^. The QM region (Figure S1) comprised 127 atoms. Specifically, the GTP, the catalytic water molecule (W_nuc_), the Mg^2+^ ion coordinated with two water molecules (W1 and W2), and the side-chain fragments of RhoA residues Lys18, Thr19, and Gln63, as well as the Arg1735 residue of RhoGAP. These residues were truncated at the Cα–Cβ bond. For residue Thr37, the backbone atoms were included, given the importance of its carbonyl group in orienting W_nuc_. Electronic structure calculations employed the Gaussian and plane-wave approach implemented in the Quickstep module, ^57^ using double-zeta plus polarization basis sets ^58^ for valence orbitals and a plane-wave cutoff of 320 Ry for the electron density. Goedecker–Teter–Hutter pseudopotentials were used to model atomic cores.^59,60^ Wavefunction optimization was achieved via the orbital transformation method with a convergence criterion of 1 × 10^−6^ for the electronic gradient.^61^ The capping of the covalent bond cut at the QM/MM boundary was handled using the hydrogen link atoms,^62,63^ whereas electrostatic embedding was carried out following the protocol of Laino *et al*. ^64^.

Simulations were performed in the NVT ensemble at 300 K using a velocity-rescaling thermostat, with a 0.5 fs integration time step over 6 ps. Convergence of the QM/MM MD simulations was assessed by monitoring RMSD, selected interatomic distances, total QM+MM potential energy, and temperature.

### QM/MM MD Metadynamics

We performed QM/MM metadynamics (MTD) simulations to assess the GTP hydrolysis mechanism in the RhoGAP:RhoA system and compute the associated free energy landscape. The rate-determining step was crossed in both forward and backward directions, verifying that the identified transition state was not an artifact of the applied bias, and convergence was ensured by performing multiple independent replicas, following an established computational protocol ^65^. During MTD simulations, Gaussian hills were added every 25 fs (corresponding to 50 MD steps). For chemical transformations, a time step of 0.5 fs was employed. The height of the deposited hills was set to 0.750 kcal·mol^−1^ for most simulations, except MTD2 and MTD4′/MTD4′′ (Table S1): MTD2 employed 3.0 kcal·mol^−1^ hills to overcome a barrier too high to be biologically relevant, with no additional replicas, whereas MTD4′/MTD4′′ used 0.250 kcal·mol^−1^ hills to avoid overfilling the low barrier and obtain accurate free-energy profiles.

The width of the Gaussian functions was defined according to the average oscillations of the collective variables (CVs) in unbiased QM/MM MD simulations ^65,66^ and are reported in Table S1.

The CVs were defined to describe the key bond-forming and bond-breaking events along the reaction coordinate. For the reaction path connecting REA to INT4 (Scheme 1), we used two CVs (CV1 and CV2 in FigureS2A). CV1 corresponds to the difference between the breaking bond (O3β:GTP– Pγ:GTP) and the forming bond (O:W_nuc_–Pγ:GTP), capturing the nucleophilic attack on the GTP γ-phosphate; and CV2, describing the proton shuttle mediated by Gln63, is defined as the difference between the distance of the transferring hydrogen from the side-chain nitrogen of Gln63 (d[Hε21– Nε2:Gln63]) and the distance of side-chain oxygen of Gln63 and the hydrogen of the nucleophilic water (d[Oε1:Gln63–H1:W_nuc_]). For the reaction path connecting REA to INT4 (Scheme 1), two independent MTD simulations were performed. The first replica (MTD1 in Table S1) included a full recrossing event, covering the entire pathway REA → INT4 → REA (Figure S2), the second MTD run (MTD1’ in Table S1) sampled instead only the forward path from REA to INT4.

For the INT3 → PROD reaction path I (Scheme S3), we used one CV (CV3, Figure S3A), defined as the distance between H1 and Nε2 of Gln63* and a single MTD simulation was carried out (MTD2, Table S1).

The INT3 → PROD reaction path II (Scheme S3) we employed two CVs (CV4 and CV5, Figure S4A). CV4 corresponds to the distance between H2:Pi and Nε2:Gln63*, while CV5 is defined as the difference between the H1–Oε1(Gln63*) distance and the H1:Gln63*–O(Pi) distance. Two independent metadynamics simulations (MTD3 and MTD3′, Table S1) were done to account for this part of the reaction mechanism.

For the INT4 → PROD reaction path III (Scheme S3), the simulations started from the structure of INT4 obtained in MTD1. An additional water molecule (W3, Figure S5) was included in the QM region, yielding a total of 130 QM atoms. Here we used two CVs (Figure S5A): CV6, defined as the H1:Gln63*–O:W3 distance, and CV7, defined as the H1:W3–Nε2:Gln63* distance, running three independent metadynamics simulations (MTD4, MTD4′ and MTD4′′ Table S1).

The variability in the calculated Helmholtz free energy barriers across different replicas was used to estimate the errors. The convergence criterion was defined as differences (ΔF) in the main minimum between independent free energy surfaces (FES) being smaller than ∼1 kcal/mol, in agreement with standard protocols for ab initio metadynamics.

Collectively we performed 71 ps of QM/MM MD/MTD simulations. Analysis was performed on all simulation replicas.

#### Analysis

Analyses were done with the CPPTRAJ package of the amber tools,^40^ MDAnalysis ^67^ and ProLIF^68^ python libraries. MEPSAnd (Minimum Energy Path Surface Analysis over n-dimensional surfaces)^69^ was used to identify the stationary points of the free energy surface. Visual Molecular Dynamics (VMD)^70^ and Chimera software^71^ were used to inspect the trajectories and to analyze crystallographic water molecules in the structures of several GTP-bound Rho GTPase/GAP complexes (PDB IDs: 1GRN, 2NGR, 1AM4, 1TX4, 1OW3, 3MSX, 5IRC).^71–74^

Water molecule tracking was performed using AQUA-DUCT v.1.0. ^75^ on the first 300 ns of the simulations to provide a representative view of water behavior. To track water molecules entering the active site via a back cavity, we defined a sphere of 6.5 Å radius centered on Arg1735 and Arg1740 of RhoGAP and Gln63 of RhoA. To assess water entry at the interface, a sphere of 4 Å radius centered on Lys12 and Gly14 of RhoA and Ser1737 and Gly1738 of RhoGAP. Path trimming was performed using AutoBarber with Scope set to the RhoGAP:RhoA complex and default parameters for all other options. Water pockets and local hot spots were identified using the pond module, and visualization was carried out in PyMOL 3.0.0. The radial distribution function of water molecules with respect to Oε1 and Nε2 of Gln63/Gln63* was calculated using the CPPTRAJ module of the AmberTools package. ^40^

Structural superpositions of the 64 human RhoGAPs were performed using BioPython v.1.86 ^76^ and AlphaFold3 ^77^ models, with pairwise alignments carried out on the full-length sequences to compare conserved structural motifs and active-site geometry.

## Results and Discussion

### RhoGAP:RhoA-GTP Forms a Stable Catalytically Competent Complex

To generate a catalytically competent structure of RhoGAP:RhoA-GTP complex we initially equilibrated the reactant (REA) structure via multiple 500 ns-long classical MD simulations (Figures S6-S7). As expected, residues of switch II exhibited reduced flexibility in the RhoGAP:RhoA complex, as compared to isolated RhoA^8,78^ (Figures S8-S9), by establishing non-covalent interactions (NCIs) at RhoGAP/RhoA-GTP interface at three main sites (Figure 1C and Table S2). For clarity, hereafter residues belonging to GAP protein will be underlined. The first anchoring site involved a3 helix of RhoA (Figure 1), where GAP’s residues <u>Arg1742</u> and <u>Arg1744</u> established persistent hydrogen (H-)bonds and salt-bridge interactions (Figure S10) with the RhoA’s residues Asp90 and Glu97, respectively, as observed in the X-ray structure.^19^ Additionally, the RhoGAP’s residues <u>Gly1738</u> and <u>Ala1740</u> residues H-bonded to Asn94:RhoA (Figure S10). The second anchoring site (Figure 1C) involved the RhoGAP’s arginine finger (<u>Arg1735</u>), which H-bonded and engaged cation– π interaction to Gln63 and Tyr34 of RhoA. Additionally, persistent interactions were established between RhoGAP’s residues <u>Arg1776</u> with RhoA’s residues Glu64 and Asp65 as well as <u>Ile1848</u> and RhoA:Tyr66. The third anchoring point (P-loop anchor, Figure 1C) involves GAP residue <u>Ser1737</u>, which transiently H-bonded to RhoA’s Gly14 of the P-loop (Figure S10).

In the RhoGAP:RhoA complex the catalytic core comprises the <u>Arg1735</u>:RhoGAP/Gln63:RhoA residue pair, along with Thr17 and Thr37:RhoA, two water molecules, and two GTP oxygens which form the Mg^2+^ ion coordination sphere (Figure 1B). Throughout the MD simulation GTP remained strongly bound to the active site, establishing several H-bonds and salt-bridge interactions, including the ones with <u>Arg1735</u> and Gln63 (Figures S11 and Table S3). In addition, the nucleophilic water (W_nuc_) H-bonded with the backbone carbonyl group of Thr37 and with the carbonyl of the Gln63 sidechain, thus being well oriented to perform the nucleophilic attack on the GTP γ-phosphate (Figure S12). In contrast to previous studies, ^79,80^ during MD simulations no additional water molecules entered the catalytic site to assist deprotonation (activation) of the nucleophilic one. A closed inspection of the crystal structures of different GTP-bound Rho GTPases/GAP supported this finding confirming that no additional waters were present nearby the nucleophilic one (see *Analysis* in the Methods section).

### Gln63 Amide to Imide Tautomerization Supports GTP Hydrolysis

We next aimed to characterize the mechanism of GTP hydrolysis in the RhoGAP:RhoA complex (Scheme 1). Starting from a representative frame of classical MD simulation, we relaxed the reactant state by performing 6 ps of unbiased QM/MM MD simulation, using the DFT-BLYP-D3 level of theory ^55,54,81^ for the QM region, and AMBER force fields for the MM part (Figure S13). The resulting equilibrated Michaelis complex (Figure S14A) maintained a stable geometry (average active site RMSD of 0.38 ± 0.05 Å relive to the crystal structure), and H-bond network around GTP. The GTP was anchored in catalytic site (Table S4) by *(i)* its coordination to the Mg^2+^ ion via its O2β:GTP and O2γ:GTP oxygen atoms (distances (d) = 2.04 ± 0.08 Å and 2.05 ± 0.08 Å, respectively) and *(ii)* H-bonds between Lys18 with GTP: (Lys18:Hε atom H-bonds to O1b (d = 2.28 ± 0.21 Å), and to O1b (d = 1.77 ± 0.16 Å), and Hε2 H-bond to O3g (d = 1.60 ± 0.12 Å). Moreover, <u>Arg1735</u> stabilizes the γ-phosphate through H-bonding to O1g (d = 1.67 ± 0.13 Å), O2A (d = 1.97 ± 0.25 Å) and O3b (d = 2.26 ± 0.36 Å). Gln63 H-bonded to the O1g atom of GTP (d = 2.05 ± 0.22 Å) well orienting it to undergo the water nucleophilic attack (Figure S14B). Likewise, the H-bonds between the nucleophilic water (W_nuc_) to the carbonyl groups of Gln63 side chain, and Thr37 backbone (Figure S14B) optimally placed W_nuc_ to perform the nucleophilic attack (i.e. the oxygen of W_nuc_ was at a distance 2.8 ± 0.4 Å from the γ-GTP phosphate (Pγ)). This H-bond network also hindered the orientation of W_nuc_ required for the solvent- or substrate-assisted pathways (Scheme S1) proposed in other studies. ^21,33^

Next, to accelerate the reaction we performed QM/MM metadynamics (MTD) simulations. MTD allows a detailed characterization of the free-energy landscape of enzymatic processes, including phosphate hydrolysis or transesterification reactions, as a function of few collective variables (CVs). ^31,82–84,65^

**Scheme 1.**
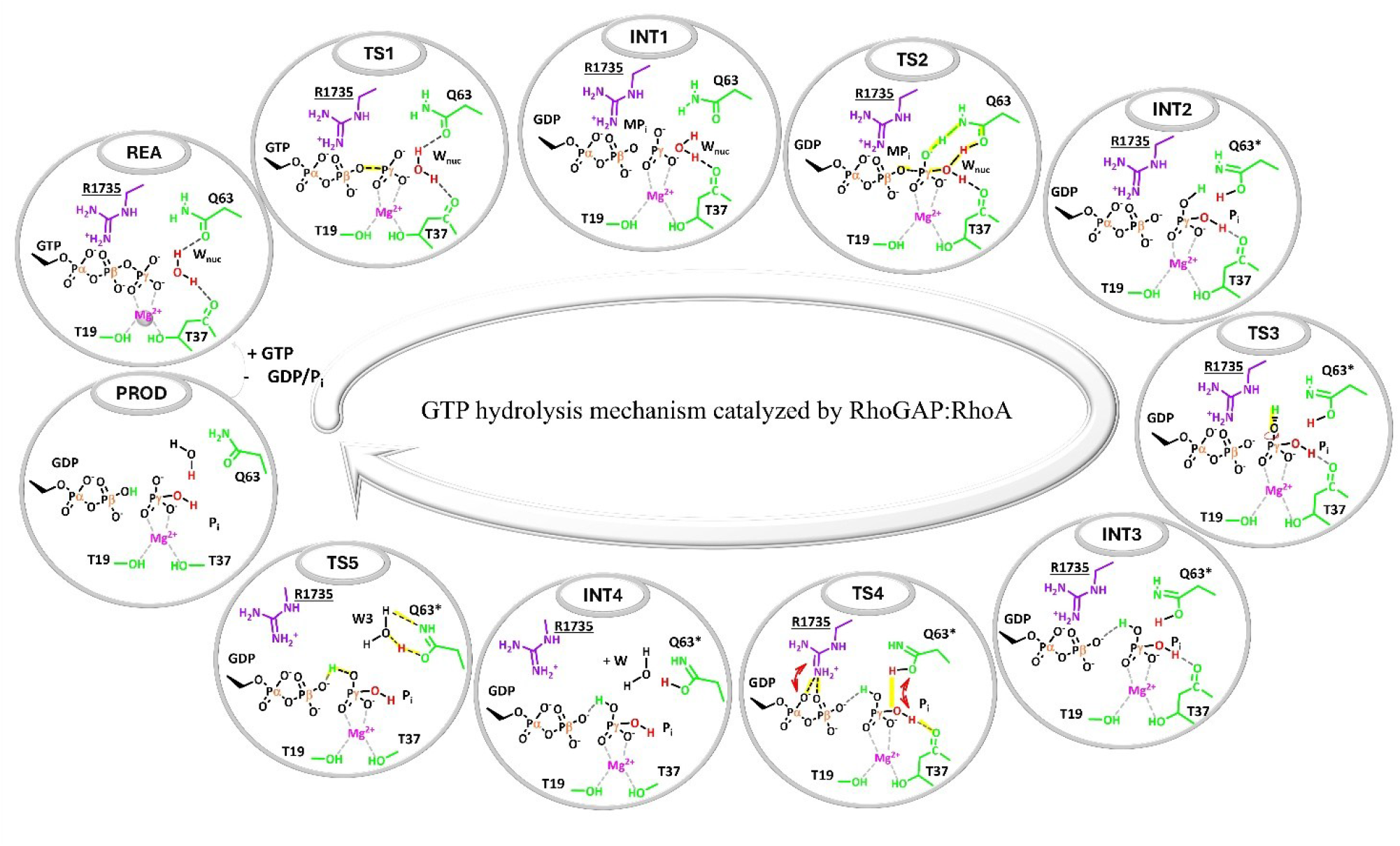
Sketch of GTP hydrolysis mechanism catalyzed by RhoA:RhoGAP complex featuring a solvent-assisted restoration of the catalytic site. The nucleophilic water molecule is in red, residues belonging to RhoGAP and RhoA are in purple and green, respectively. Residues are shown only partially for clarity.

To this end, we used two CVs (Figure S2): CV1 was defined as the difference of the distances between the breaking (d1 = O3β:GTP–Pγ:GTP) and the forming (d2 = O:W_nuc_–Pγ:GTP) bonds of the nucleophilic substitution reaction while CV2 was defined as the difference between the distance of the transferring hydrogen from the nitrogen of Gln63 side chain to the γ-phosphate (d3 = Hε21– Nε2:Gln63) and the distance between the oxygen of the Gln63 side chain and the hydrogen (H1) of the nucleophilic water (d4 = Oε1:Gln63–H1:W_nuc_). The CV2 captures the proton shuttle from the nucleophilic water to the GTP oxygen via Gln63, mediated by its tautomerization. While previous studies have proposed that Gln63 could act as a general base, ^85,86^ the high pK_a_ value of the same Gln residues in Ras ^87^ suggests that an amide→imide tautomerization mechanism may be a plausible route.

In MTD simulations GTP hydrolysis starts (REA → INT1) with the nucleophilic attack of W_nuc_ on the γ-phosphorus atom of GTP (Pγ:GTP), which, by overcoming an Helmholtz free energy barrier (ΔF^‡^ TS1, Figure 2) = 10.5 ± 1.2 kcal·mol^−1^, leads to the formation of a metastable intermediate (INT1, Figures 2), ΔF = 9.7 ± 0.5 kcal mol^-1^, consistently with a dissociative mechanism. Next, by overcoming a further ΔF^‡^ = 8.0 ± 0.7 (TS2 in Figure 2) the formation of H_2_PO_4_^−^ (Pi), and GDP occurs concertedly with proton transfer from the nucleophilic water (W_nuc_) to the carbonyl side chain oxygen of Gln63, which in turn donates its amide proton to the departing γ-phosphate. Thus the nucleophilic attack is coupled to an amide-to-imide tautomerization of Gln63 (hereafter denoted Gln63*) and, collectively, the formation of the the first stable intermediate (INT2) requires overcoming a ΔF^‡^ = 17.7 ± 0.7, which aligns well with the experimentally measured rate constant for RhoGAP-catalyzed GTP hydrolysis (k_cat_ ≈ 60 min^−1^). ^16,19^ The free energy surface and time evolution of the CVs for all catalytic steps are shown in Figure S2.

Similarly to other hydrolases^88–90^ GTP hydrolysis then proceeded (INT2 → INT3) through an H-bond rearrangement in which the H_2_PO_4_^−^ hydrogen, newly transferred from Gln63, reoriented to form an intramolecular H-bond with O3β:GDP (O3β:GDP–Hε21:Pi = 1.48 ± 0.13 Å). This rearrangement— which requires the O3β:GDP–Pγ:Pi–O1γ:Pi–Hε21:Gln63 dihedral to rotate from –137.9 ± 12.7° in INT2 to –3.36 ± 6.1° in INT3 (Figure S13)—proceeded surpassing a barrier of ΔF^‡^ = 7.9 ± 0.6 kcal·mol^−1^ (Figure 2), yielding INT3 at ΔF = 13.7 ± 0.8 kcal·mol^−1^.

The next step of the catalytic process (INT3 → INT4, Figure 3A) required a further weakening of the interactions between H_2_PO_4_^−^ and the catalytic site. Here the H-bond between the Pi hydroxyl group (–OH, deriving from the original W_nuc_) and the backbone of the Thr37 (Figure S14) broke, concurrently with a weakening of arginine finger H-bond to the GDP. Specifically, the distances between HH21:<u>Arg1735</u> and O3β:GDP (d4), HH22:<u>Arg1735</u> and O2α:GDP (d5), and Hε:<u>Arg1735</u> and O1γ:GDP (d6)—which measure 1.65 ± 0.19 Å, 2.05 ± 0.19 Å, and 2.09 ± 0.12 Å, respectively, in INT3—increase to 2.31 ± 0.14 Å, 3.05 ± 0.10 Å, and 2.71 ± 0.10 Å in INT4 (Figure S14). This transition required crossing a ΔF^‡^ = 7.3 ± 0.7 kcal·mol^−1^ and was endothermic by ΔF = 17.7 ± 0.3 kcal·mol^−1^.

**Figure 3.**
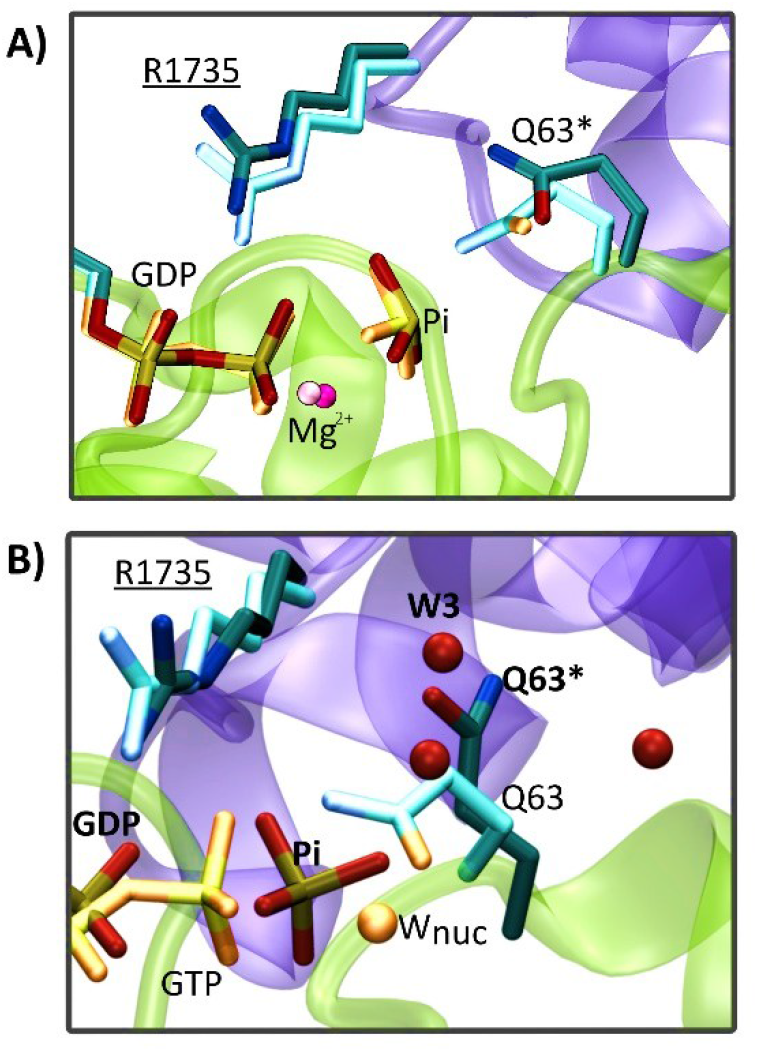
A) Superposition of INT3 and INT4 structures on the Helmholtz free energy surface shown in Figure 2. Residues are displayed as sticks representation, using light and bright colors for INT3 and dark and intense colors for INT4. B) Superposition of the structure of REA and INT4 obtained from Classical MD. Residues are shown as sticks representation, with light and bright colors for REA and dark and intense colors for INT4. Oxygen atoms of water molecules located within a 2.5 Å of Q63 (light red for the REA state) and Q63* (intense red for INT4) are displayed. For clarity, the cartoon representation is shown only for INT3 in panel (A) and for the REA state in panel (B).

### Glutamine Imine Tautomer Loosens the RhoGAP–RhoA Complex, Enabling Water Entry and Active-Site Restoration

Restoring the catalytic site requires the Gln63 imide tautomer to revert to its amide form. We assessed three potential recovery pathways (Scheme S3) and calculated their free-energy profiles to identify the most favourable route.

### Intramolecular Gln63 proton transfer (Path I)

We initially attempted to restore the standard Gln63 amine form by enforcing an intra-Gln63 proton transfer. QM/MM metadynamics simulations starting from a representative structure of INT3 and using as using as CV the distance between Gln63*:H1 and Nε2:Gln63* (CV3, Figure S3) revealed that this path (Path I) was not energetically viable since it required overcoming a ΔF^‡^ of about 35 kcal·mol^−1^ (Figure S3).

### Phosphate assisted Gln63 restoration (Path II)

We next explored a pathway in which the dissociated g-phosphate assists the proton transfer from Oε1:Gln63* to Nε2:Gln63*. Starting from a representative INT3 structure we performed a metadynamic simulations using two CVs (CV4 and CV5 in Figure S4), defined as the distance between H2:Pi and Nε2:Gln63*, and the difference of the distances H1-Oε1 of Gln63* and H1: Gln63*-O:Pi. These CVs account for the proton transfer from the imide Gln63* Oe: to its Ne, mediated by the –OH group of the g-phosphate, while retaining the rotational freedom of the γ-phosphate, known to be important in Ras:GAP. ^32^ In these simulations the phosphate aided the proton transfer from Oε1:Gln63* to Nε2:Gln63* by forming a three-centre transition state where the proton simultaneously interacted with O:Pi, Oε1:Gln63* and Nε2:Gln63* (Figure S4D). PROD formation occurred with ΔF^‡^=11.8 kcal·mol^−1^ and ΔF= –2.5 kcal·mol^−1^ (Figure S4). Interestingly, during PROD relaxation via 6 ps of QM/MM MD we observed rearrangements of the H-bond network. While <u>Arg1735</u> finger stably H-bonded to the diphosphate (HH21:Arg1735 and O3β:GDP) (Figure S14 and Table S4), the GDP and Pi remained bound to the Mg^2+^ ion and RhoA P-loop residues, the interactions at the RhoGAP:RhoA interface started loosening. Thus, to further investigate the evolution of RhoGAP:RhoA interfacial contacts, we performed multi-replica 500 ns long classical MD simulations of PROD. These revealed an additional reduction in interfacial interactions (Figure S10) and a progressive structural destabilization of the complex (Figure S6).

### Water assisted Gln63 restoration (Path III)

We finally explored a previously unexamined pathway where Gln63 restoration is assisted by water (Path III, Scheme S3). During the catalytic process (Figure 2), the transition from INT3 to INT4 induces a more open active site conformation, in which Gln63* is more solvent exposed rather than interacting with the Pi (Figure 3A).

Interestingly, MD simulations of INT4 revealed a rearrangement of the H-bond network of RhoGAP’s residue <u>Ser1737</u>, which, while H-bonding with the backbone of Gly14 in REA, and formed a new H-bond to Asp13:RhoA side chain in INT4 (Figure S10). As a result of this H-bond remodeling, the P-loop anchor site weakens (Figure 3), inducing decreased interfacial interactions, and a structural destabilization of RhoGAP:RhoA (Figures S6–S7) complex. The loosening of interfacial H-bond network allowed the rapid entry of water molecules during classical MD simulations. Indeed, water tracking and Radial Distribution Function (RDF) analyses revealed the entry of 1–3 water molecules into the RhoGAP:RhoA interface, depending on the replica (Figures 3B, S17–S18), with one molecule consistently positioned in proximity to Gln63* imide group. This suggests that solvent may likely mediates Gln63 reverse imide-to-amide tautomerization.

To explore this intriguing possibility, we performed three independent QM/MM MTD simulations (MTD4, MTD4′, and MTD4″; Table S1, for a cumulative sampling time of 11 ps) with an additional water molecule (W3, Figure S5A) in the catalytic site. This water molecule was introduced in the QM region to mimic the behavior of the W3 during the MD simulations (Figure 3B). Here we used two CVs to enhance the solvent-mediated proton transfer from Oε1 → Nε2 of Gln63. CV6 was defined as the distance between Hε1 bound to Oε1 of Gln63* and the oxygen atom of W3, describing the proton transfer from the Gln63 Oe1 to W3 oxygen, whereas CV7 was defined as the distance between H2 of W3 and Nε2 of Gln63*, thereby describing the subsequent proton transfer from W3 to the imide nitrogen Ne. In all MTD simulations, W3 promoted the proton transfer from Oε1:Gln63* to Nε2:Gln63* by forming a three-center transition state where the migrating proton simultaneously interacted with O:W3, Oε1:Gln63*, and Nε2:Gln63* (Figure S5D). Next, PROD formation (Figure 2) proceeded by overcoming a small free-energy barrier (ΔF^‡^ = 3.6 kcal·mol^−1^) and the process was exergonic (ΔF = –7.9 kcal·mol^−1^). In the product state, the previously transferred proton (Hε21) interacts with GDP (Figures 2 and S5). PROD is further stabilized by an extended H-bond network involving Gln63, Pi, and W3 (Figure S5).

To assess the generality of water-assisted tautomerization, we also examined if this could occur in RhoA in aqueous solution, namely after the dissociation of RhoGAP has already occurred. During 5-ps-long QM/MM MD simulations we observed a spontaneous and barrierless reverse imide-to-amide tautomerization of Gln63, with a solvent molecule acting as a proton shuttle between Oε1 and Nε2, consistent with the concerted water-assisted tautomerization mechanism reported for isolated amino acids in water solution. ^91,92^

These findings suggest that the water-assisted mechanism for Gln63 recovery is the lowest-energy pathway occurring at INT4 or in isolated RhoA after the RhoA/RhoGAP complex has dissociated.

### Catalytic and RhoGAP–RhoA Contact Determinants are Conserved Across the Human RhoGAP Family

To determine whether the catalytic mechanism uncovered here is shared across other Rho GTPases, we carried out integrated bioinformatic and structural analyses of human RhoGAPs. This protein family comprises 66 members involved in diverse signaling pathways, whose functional specificity is governed by regulatory domains distinct from the catalytic core. ^93^ Out of these, 56 contain a conserved catalytic domain that terminates Rho signaling by accelerating GTP hydrolysis. ^93^

Since our analyses showed that rearrangement of the H-bond network at the RhoGAP:RhoA interface promotes water entry at INT4 (Figure 3B), thereby weakening interfacial contacts and allowing the entry of solvent, we inspected which residues contribute to this process. In addition to the RhoGAP conserved arginine finger (<u>Arg1735)</u> and RhoA catalytic Gln63 residue, both <u>Ser1737:RhoGAP</u> and Asp13:RhoA actively participate in this remodeling. We also identified persistent interactions between RhoGAP residues <u>Gly1738</u>, <u>Ala1740</u>, <u>Arg1742</u>, <u>Lys1772</u>, and <u>Ile1848</u> and RhoA residues Tyr34, Glu64, Asp65, Tyr66, Arg68, Glu93, and Asn94 (Figures S10 and S16). Sequence analysis of Rho GTPases revealed these residues are largely conserved across the family (Figure S19). Further sequence analysis of the 57 human RhoGAP catalytic domains (Figure S20) revealed that <u>Arg1735</u>, <u>Ser1737</u>, and <u>Arg1772</u>—three residues that form persistent contacts with RhoA—are conserved in most RhoGAPs (57%), indicating preservation of the Rho GTPAse/GAP interfacial motif. Accordingly, for the 31 RhoGAPs carrying Arg1735, Ser1737, and Arg1772 signature, we generated structural models of their Rho-RhoGAP complexes using AlphaFold3.^77^ The resulting models (Figure S21) revealed a remarkable conservation of the catalytic geometry observed in RhoGAP:RhoA complex, as well as of relative orientation of <u>Ser1737</u>, Asp13, and of the remaining interacting residues at the Rho:GAP interface. Together, these features indicate that loosening of the RhoA:RhoGAP interface at INT4, followed by the water-assisted reverse tautomerization of Gln63, is likely a broadly conserved mechanism among many human GAPs.

## Conclusions

Small Rho GTPases hydrolyse GTP in cooperation with GAPs, but the underlying catalytic mechanism has remained debated, largely because the Rho GTPases active site lacks an obvious general base to promote GTP hydrolysis. Here we resolved the catalytic mechanism of RhoGAP:RhoA, revealing revealed that GTP hydrolysis proceeds in multiple steps with the rate limiting one being the nucleophilic attach of a water molecule to the γ-phosphate of the GTP. The computed activation free energy barrier of this step (ΔF^‡^=17.7 kcal·mol^−1^) is in excellent agreement with experimental kinetic data^16,19^ supporting the relevance of our findings. Key to this mechanism is the amide-to-imide tautomerization of Gln63, which enables a Gln63-mediated proton shuttle from the nucleophilic water to the GTP γ-phosphate. Additionally, the imide tautomer of Gln63 triggers the weakening of RhoA/RhoGAP interfacial interactions, thereby promoting solvent access to the active site, and enabling a water-assisted imide-to-amide reverse tautomerization of Gln63 to restore a catalytically competent active site.

Structural and bioinformatic analysis of human RhoGAPs indicate that the RhoGAP residue <u>Ser1173</u>, involved in the interfacial interaction remodeling with Asp13:RhoA, is conserved among most RhoGAPs, suggesting that this mechanism may be broadly shared across the family.

## Acknowledgments

AP thanks Italian Association for Cancer Research (AIRC) for the Fellowship for Italy Post-Doc. AM thanks the Italian Association for Cancer Research (AIRC) (IG grant 24514). AP, RR and AM thank CINECA, the Italian Supercomputing Center, for computational resources accessible via the ISCRA “<u>IsB32_RacMR</u>” HPC project. AP thanks “PRP@CERIC – Pathogen Readiness Platform for CERIC-ERIC Upgrade” funded by the European Union through the National Recovery and Resilience Plan (NRRP), part of Next Generation EU, as part of Mission 4 “Education and Research”, Component 2 “From Research to Business”, Investment Line 3.1 “Fund for the creation of an integrated system of research and innovation infrastructures”.

## Table of Contents

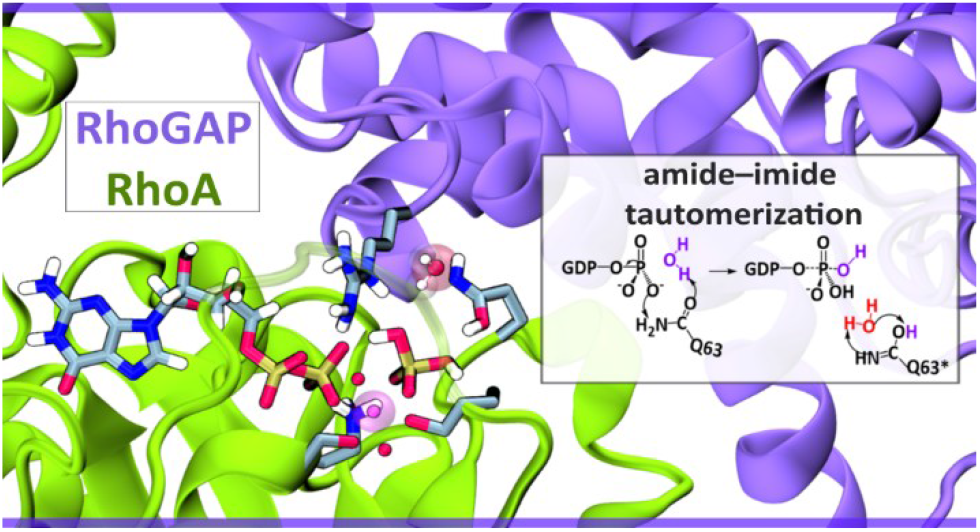

