## Supplementary material for "Glutamine Tautomerization Drives RhoGAP‑Aided GTP Hydrolysis in Small Rho GTPases": SI_AP

[a] CNR - Istituto Officina dei Materiali (IOM) c/o SISSA via Bonomea 265, 34136, Trieste, Italy.

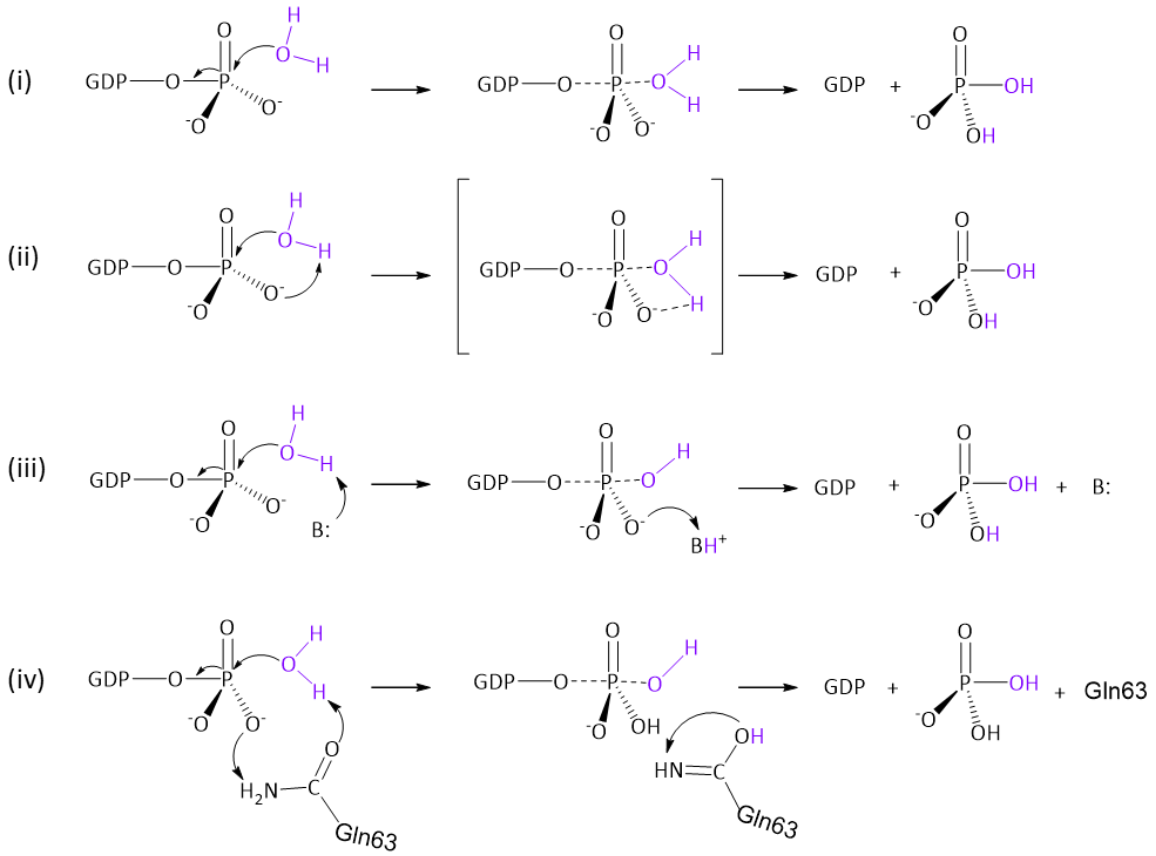


**Scheme S1.** Proposed Mechanisms for GTP hydrolysis catalysed by Rho GTPase proteins: (i) solvent-assisted mechanism; (ii) substrate-assisted mechanism; (iii) base-assisted mechanism, in which the base (B) can be either Gln or a water molecule (the latter referred to as the two-water mechanism); and (iv) amide–imide tautomerization mechanism.

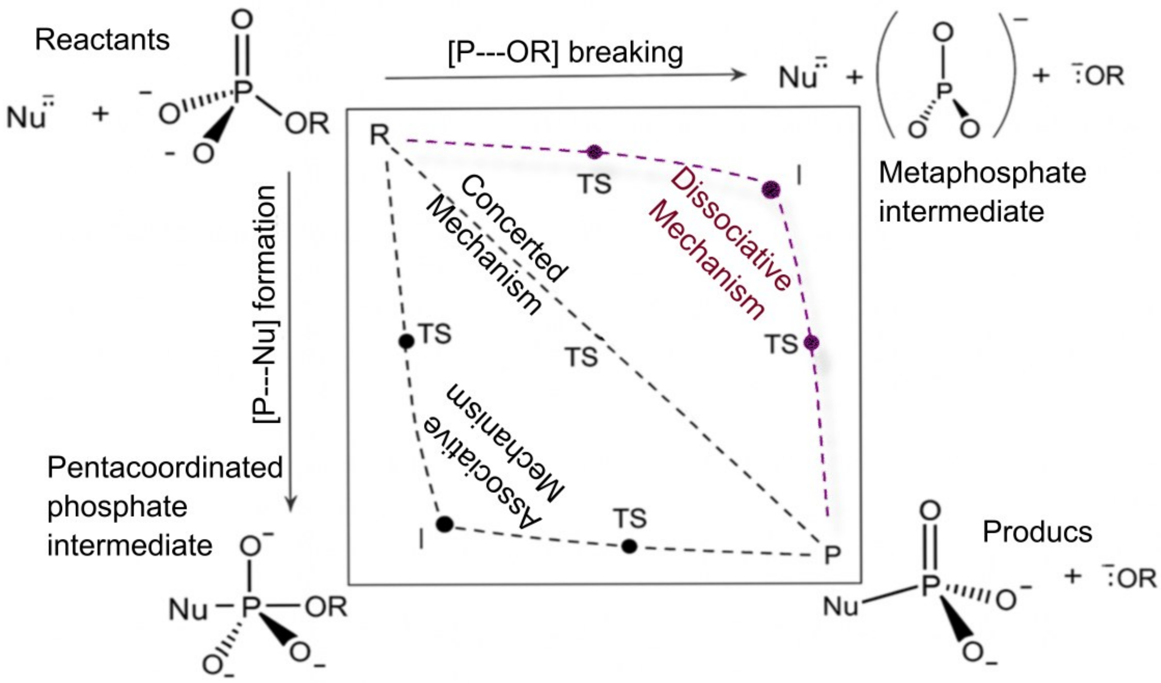


**Scheme S2.** More–O'Ferrall–Jencks representation of the general plausible reaction pathways for phosphate lysis. Three mechanisms are considered: *(i)* dissociative mechanism: proceeds via an $S_N1$ pathway in which the leaving group first dissociates from the phosphorus atom, generating a metaphosphate intermediate that is later attacked by the nucleophile (here a water molecule). *(ii)* Associative mechanism: involves the formation of a pentacoordinate phosphorus intermediate, which then releases the leaving group to generate the products. *(iii)* Concerted mechanism: proceeds through a single transition state in which bond cleavage to the leaving group and bond formation with the nucleophile occur. This mechanism may be synchronous or asynchronous but always features a single continuous transition state. The dotted lines trace the reaction pathways associated with each mechanism, described in terms of changes in the P–OR (horizontal axis) and P–Nu (vertical axis) distances. The reaction pathway by which RhoGAP:RhoA hydrolyzed GTP proceeds is indicated in purple.

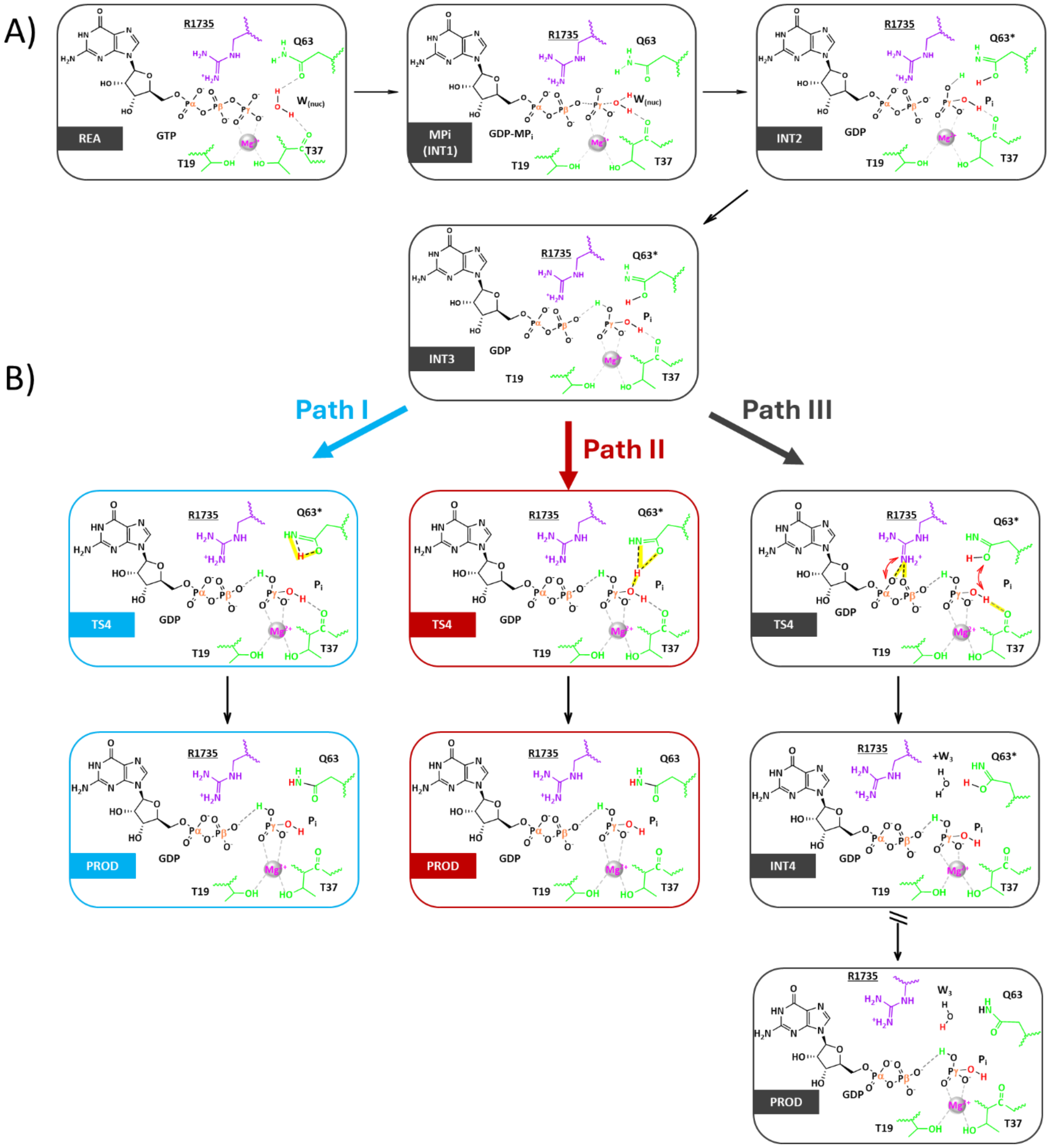


**Scheme S3.** Proposed mechanisms of GTP hydrolysis catalyzed by the RhoGAP:RhoA complex. A) Reaction pathway from leading from the reactant (REA) state to the formation of GDP and inorganic phosphate (Pi) (INT2), followed by subsequent hydrogen-bonding remodeling (INT3). B) Reverse imide to amide tautomerization of Gln63* proceeding through three possible alternative pathways. In Path I (cyan), restoration of the Gln amide occurs via a direct intramolecular proton transfer within Gln63* (i.e. direct intramolecular path). In Path II (red), starting from INT3 intermediate, proton H1ε of Gln63* is transferred from Oε to Nε2 through the phosphate (i.e. phosphate mediated mechanism). In Path III (black), progressive weakening of the hydrogen-bond interactions between the arginine finger and the phosphate groups triggers Gln63* (INT4) remodeling/opening, enabling the entry of a water molecule (W3); in this case, proton Hε1 is transferred to Nε2 tough water (i.e. water-mediated mechanism).

For clarity, only partial representations of key residues are shown.

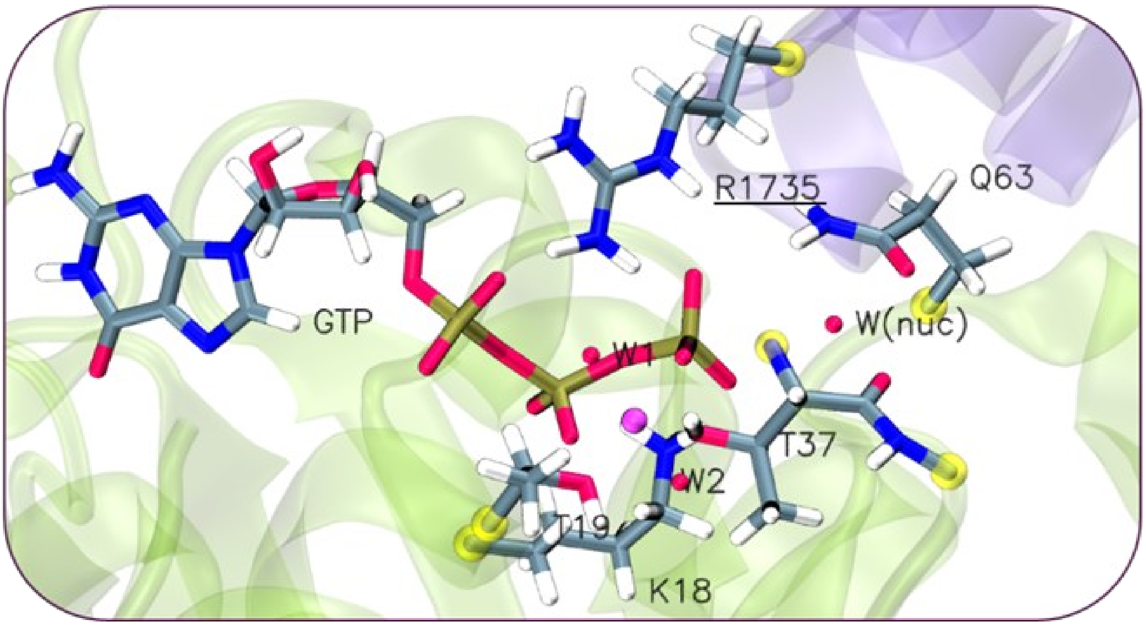


**Figure S1.** QM region composed of 127 atoms, including 6 hydrogen link atoms (yellow spheres). For clarity, hydrogen atoms of water molecules are not shown. Protein residues and the GTP are shown in sticks. Carbon, oxygen, nitrogen, phosphorous and hydrogen atoms are colored in gray, red, blue, brown, and white, respectively. $Mg^{2+}$ ion is shown as a magenta van der Waals sphere. RhoA and RhoGAP are shown as transparent violet and green new cartoons, respectively.

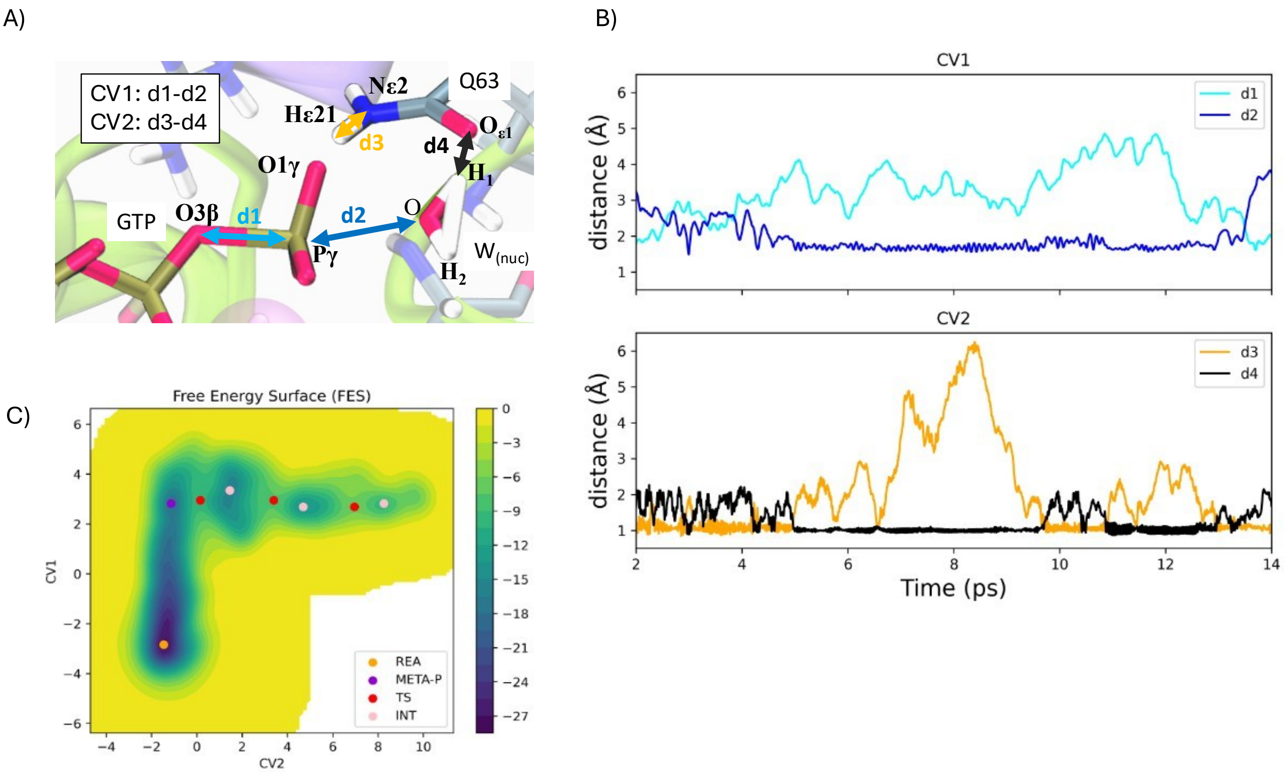


**Figure S2.** A) Definition of the collective variables (CVs) used to perform metadynamics simulations of the GTP hydrolysis reaction. CV1 corresponds to the difference between the breaking bond (d1= O3β:GTP–Pγ:GTP, cyan arrow) and the forming bond (d2= O:$W_{nuc}$–Pγ:GTP, blue arrow), capturing the nucleophilic attack on the GTP γ-phosphate. CV2, describing the proton shuttle mediated by Gln63, is defined as the difference between the distance of the transferring hydrogen from the side-chain Nε of Gln63 to the γ-phosphate (d3= Hε21–Nε2:Gln63, orange arrow) and the distance between the side-chain Oε of Gln63 and the hydrogen of the nucleophilic water (d4= Oε1:Gln63–H1:Wnuc, black arrow). B) Time evolution of the CV distances along the metadynamics trajectory. Colour palette corresponds to the arrows in panel (A). C) Free-energy surface (FES, kcal·$mol^{-1}$) projected along the CVs (atomic units). The FES reports key intermediates and transition states of the GTP hydrolysis reaction.

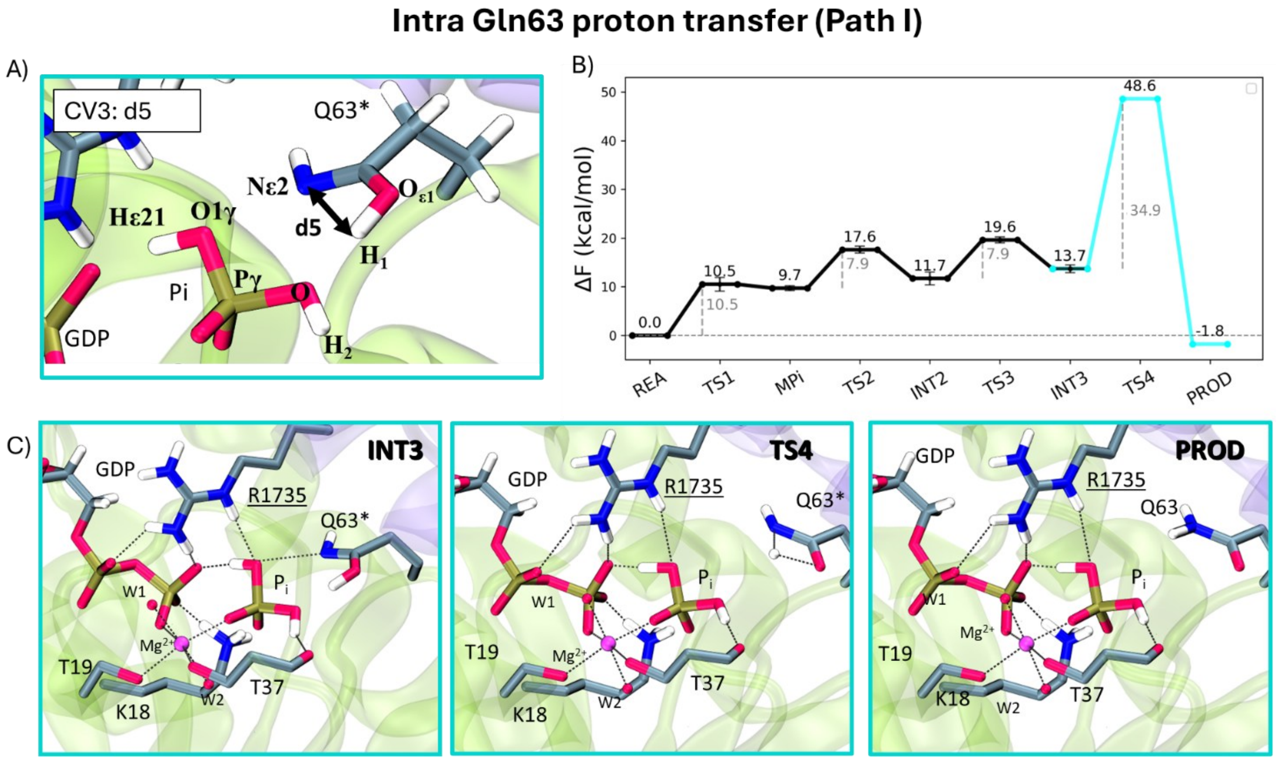


**Figure S3.** A) Definition of the collective variable (CV3) used to monitor the imide to amide reaction thought direct intramolecular Gln63 proton transfer (Path I). CV3 corresponds to distance (d5) between H1 and Nε2 of Gln63*. B) Schematic free-energy profile of the imide to amine reaction obtained at the QM(BLYP-D3(DZVP))/MM level metadynamics (in cyan). Reaction free energies (ΔF in kcal·mol$^{-1}$) and activation barriers free energy barriers are reported relative to the INT3 (Figure 2 in the main text). C) Representative snapshots of INT3, TS4, and PROD met along Path I. Protein residues and the GTP are shown in sticks. Carbon, oxygen, nitrogen, phosphorous and hydrogen atoms are colored in gray, red, blue, brown, and white, respectively. $Mg^{2+}$ ion is shown as a magenta van der Waals sphere.

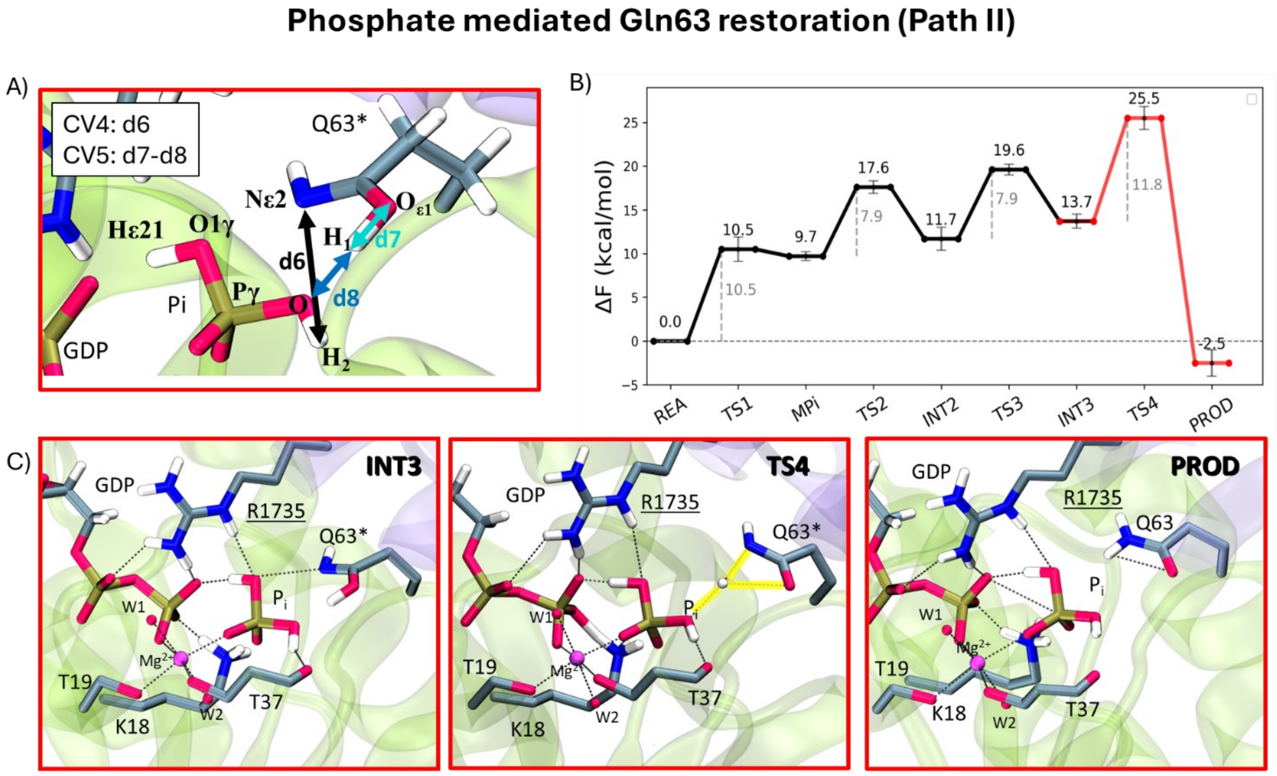


**Figure S4.** A) Definition of the collective variables *(CVs)* used to monitor the imide-to-amide tautomerization reaction assisted by Pi *(phosphate assisted path)*. CV4 corresponds to distance (d6) between H2:Pi and Nε2:Gln63*, and CV5 is the difference between the distance (d7) H1-Oε1 of Gln63* and the distance (d8) H1:Gln63*-O:Pi. B) Schematic free-energy profile of the imide-to-amide tautomerization reaction obtained at the QM(BLYP-D3(DZVP))/MM level metadynamics (red line). Reaction free energies (ΔF, kcal·mol$^{-1}$) and activation *free energy* barriers are reported relative to the INT3 *(*Figure 2*,* maim text*)*. C) Representative snapshots of INT3, TS4 and PROD along the phosphate assisted pathway. Protein residues and the GTP are shown in sticks. Carbon, oxygen, nitrogen, phosphorous and hydrogen atoms are colored in gray, red, blue, brown, and white, respectively. $Mg^{2+}$ ion is shown as a magenta van der Waals sphere.

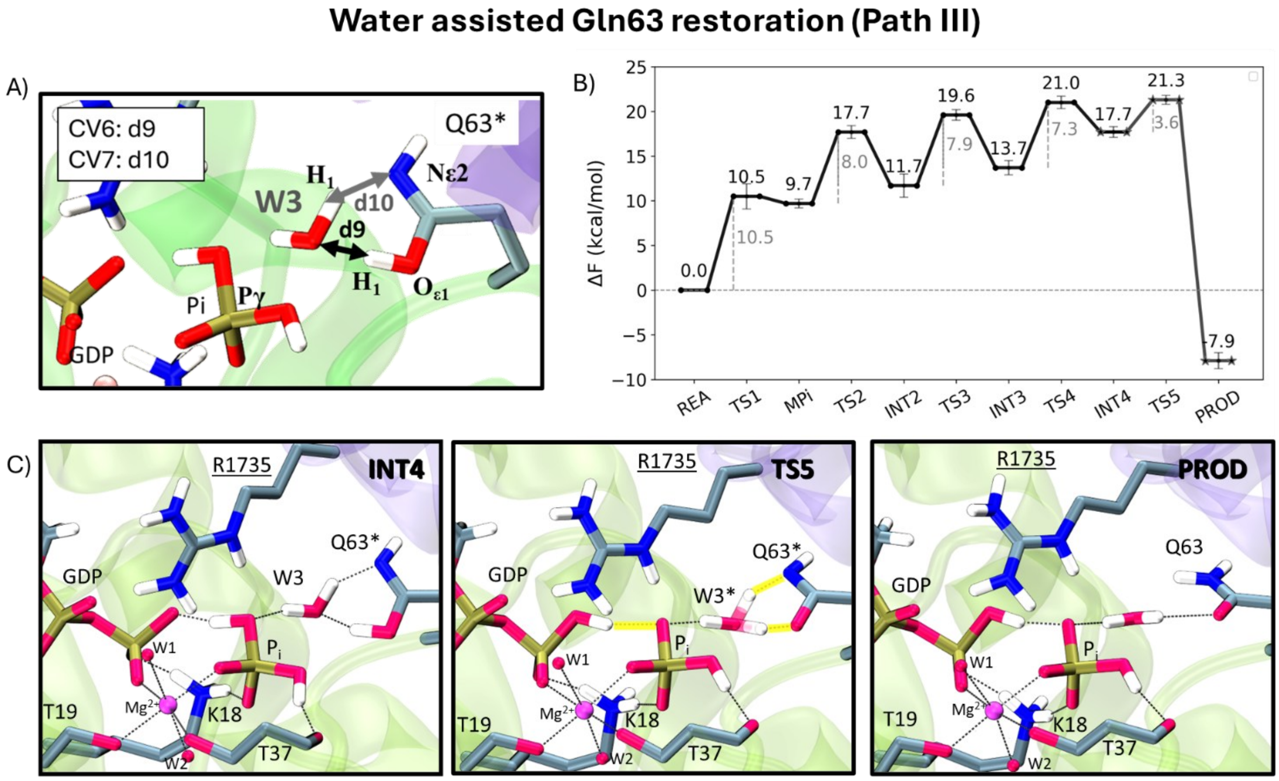


**Figure S5.** A) Definition of the collective variables *(CVs)* used to monitor the imide to amide reaction assisted by a water molecule ($W_3$) *(solvent assisted pathway)*. CV6 corresponds to distance (d9) between H1-Oε1 of Gln63* and O of W3, and CV7 corresponds to the distance (d10) between H2:W3 and Nε2:Gln63*. B) Schematic free-energy profile of the imide to amide reaction obtained at the QM(BLYP-D3(DZVP))/MM level metadynamics (states marked with star symbols). Reaction free energies (ΔF in kcal·mol$^{-1}$) and activation free energy barriers are reported relative to INT4 *(*Figure 2*,* maim text*)*. C) Representative snapshots of INT4, TS5 and PROD met along the solvent assisted pathway. Protein residues and the GTP are shown in sticks. Carbon, oxygen, nitrogen, phosphorous and hydrogen atoms are colored in gray, red, blue, brown, and white, respectively. $Mg^{2+}$ ion is shown as a magenta van der Waals sphere.

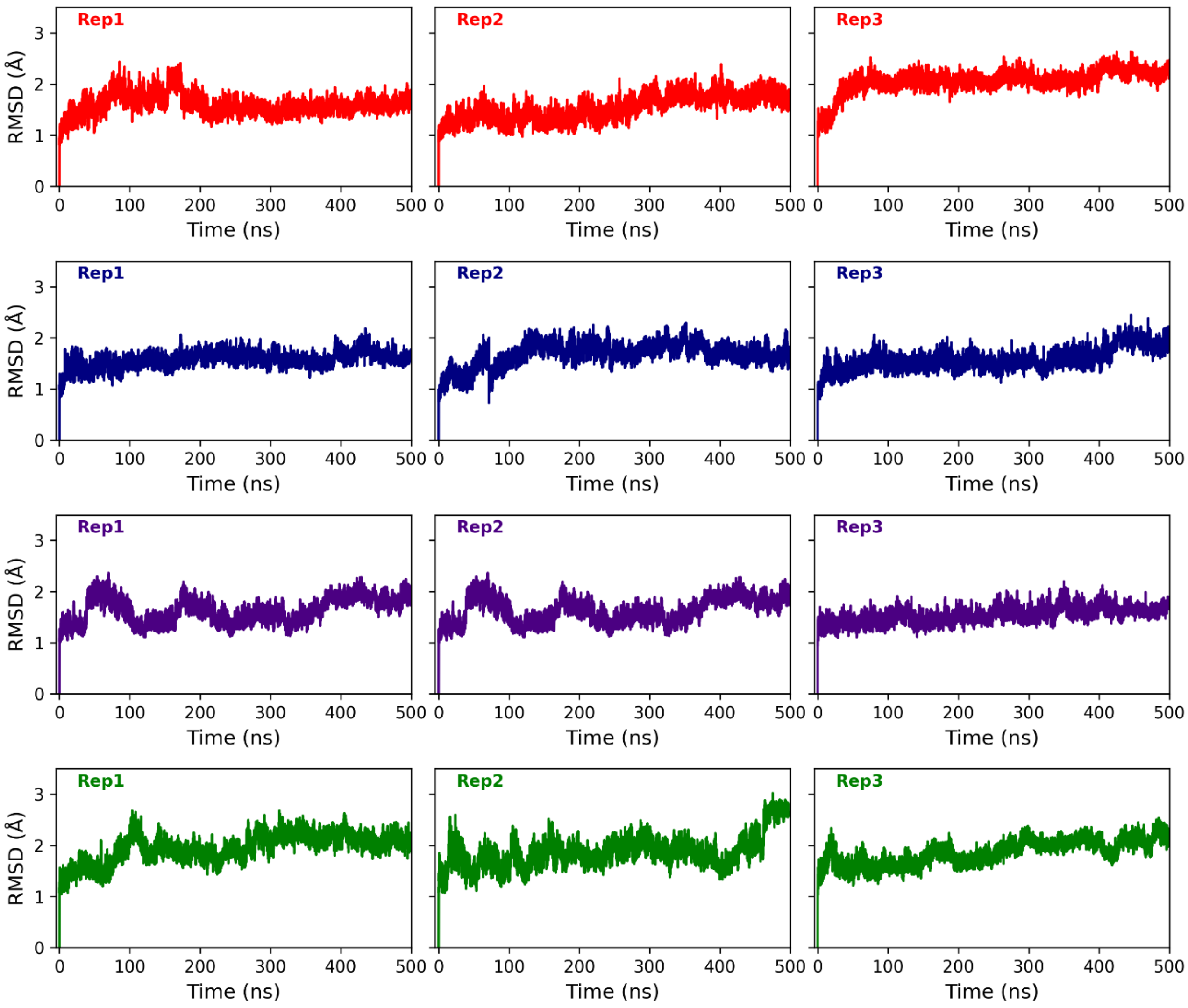


**Figure S6.** Root Mean Square Deviation (RMSD, Å) of protein backbone atoms (Cα, C, N, O) vs simulation time (500 ns) for three replicas of classical Molecular Dynamics simulations of systems a) REA (red), b) INT3 (blue), c) INT4 (violet), and d) PROD (green), defined in Scheme 1 of the main text.

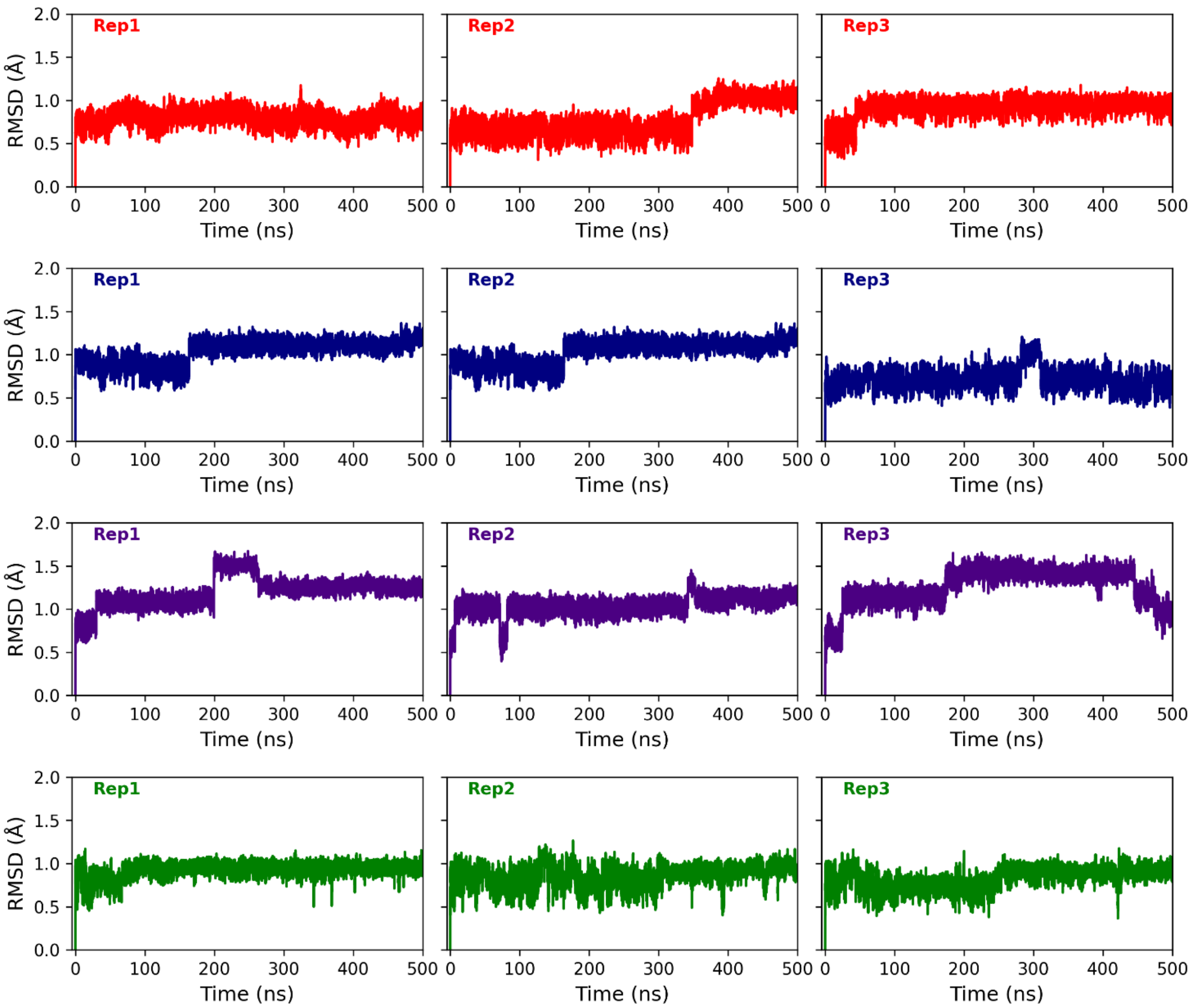


**Figure S7.** Root Mean Square Deviation (RMSD, Å) of active site residues (**Figure S1**) vs simulation time (500 ns) for three replicas of classical MD simulations of systems a) REA (red), b) INT3 (blue), c) INT4 (violet), and d) PROD (green).

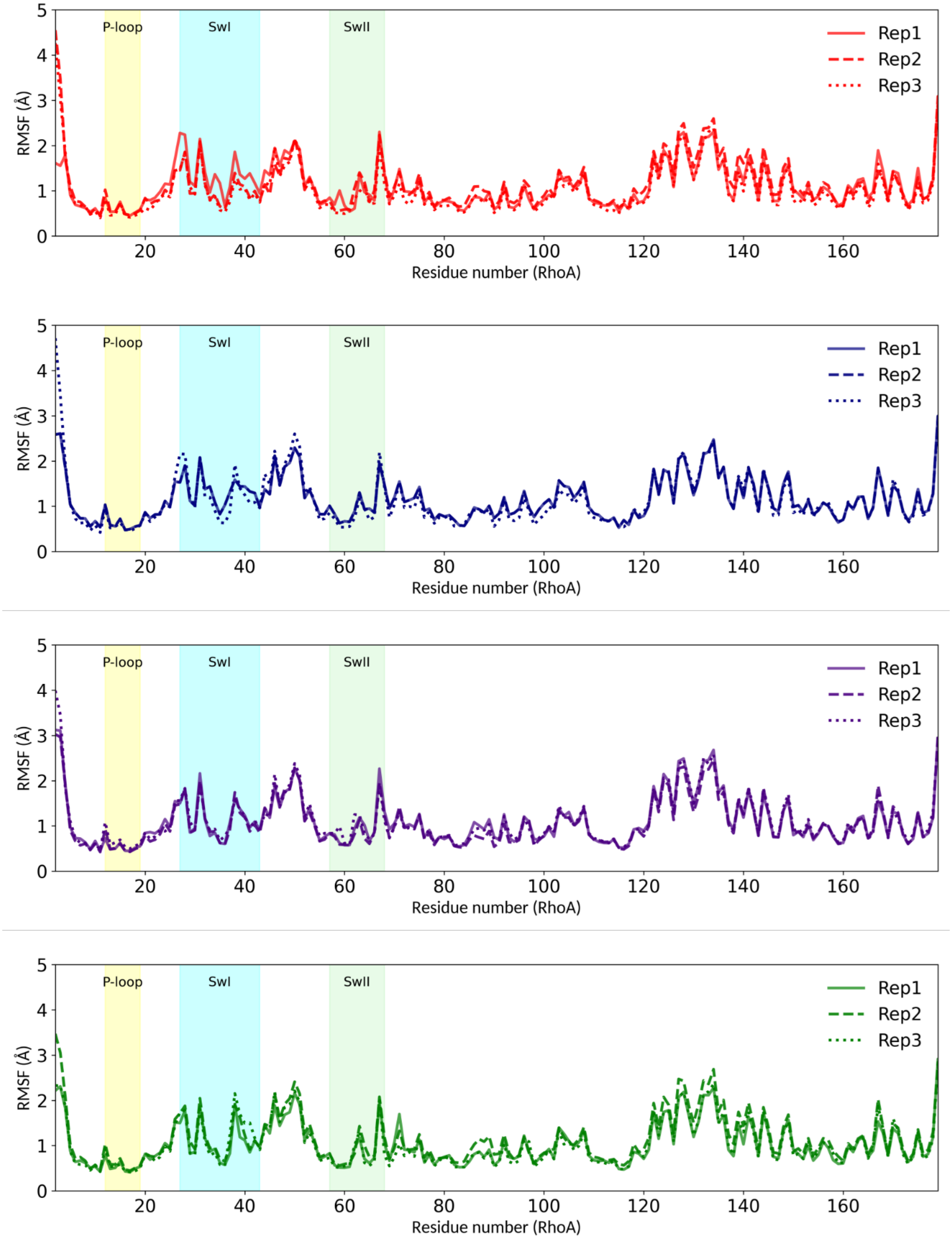


**Figure S8.** Root Mean Square Fluctuations (RMSF, Å) versus residue number. RMSF is calculated on heavy atoms of RhoA protein's residues over three 500 ns-long replicas of classical MD simulations of systems REA (red), INT3 (blue), INT4 (violet), and PROD (green). The main functional sites p-loop, switch I (swI), switch II (swII) are highlighted in yellow, light-blue, and green, respectively.

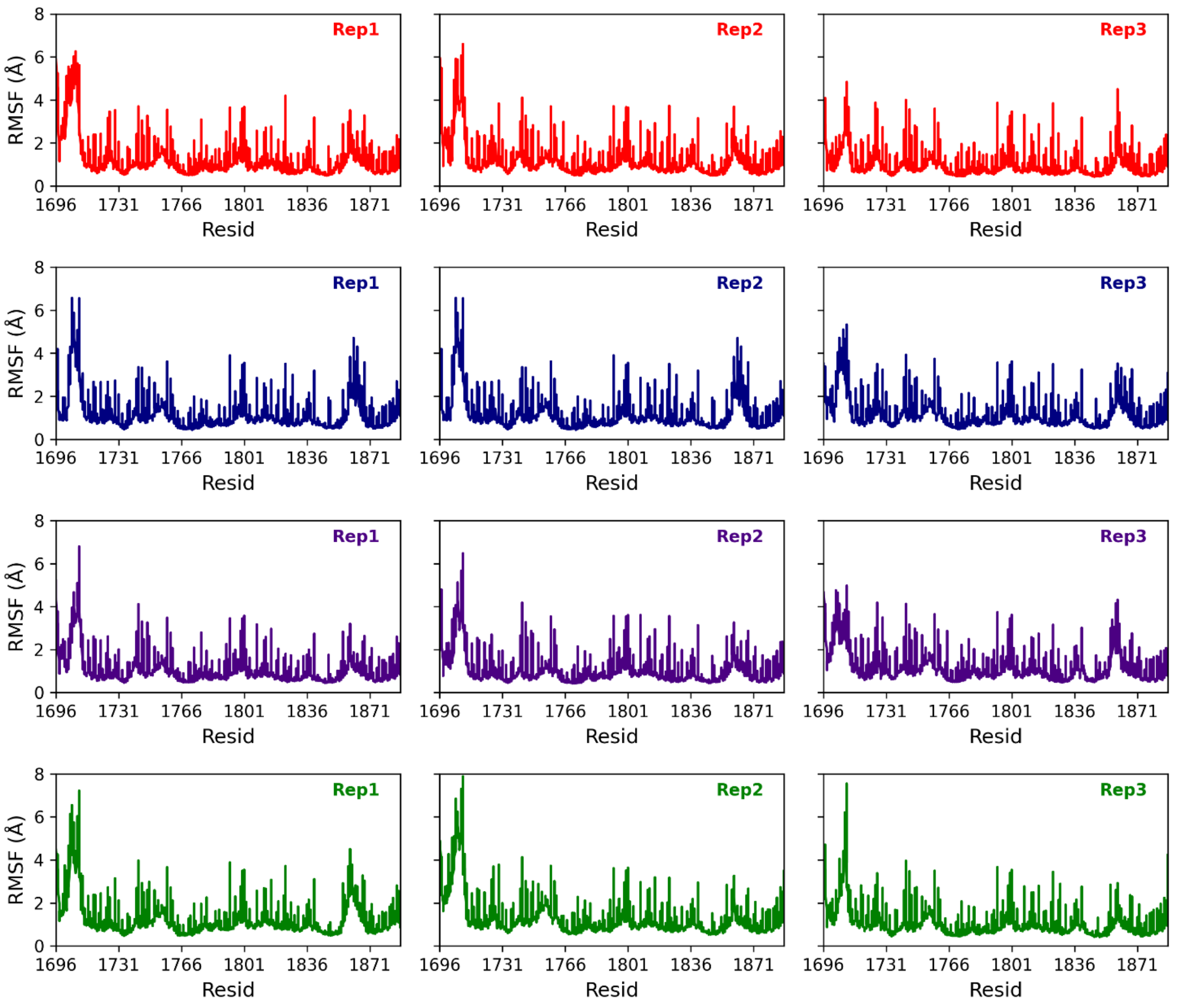


**Figure S9.** Root Mean Square Fluctuations (RMSF, Å) versus residue number calculated on heavy atoms of RhoGAP protein's residues calculated over three 500 ns-long classical MD simulations replicas of systems REA (red), INT3 (blue), INT4 (violet), and PROD (green).

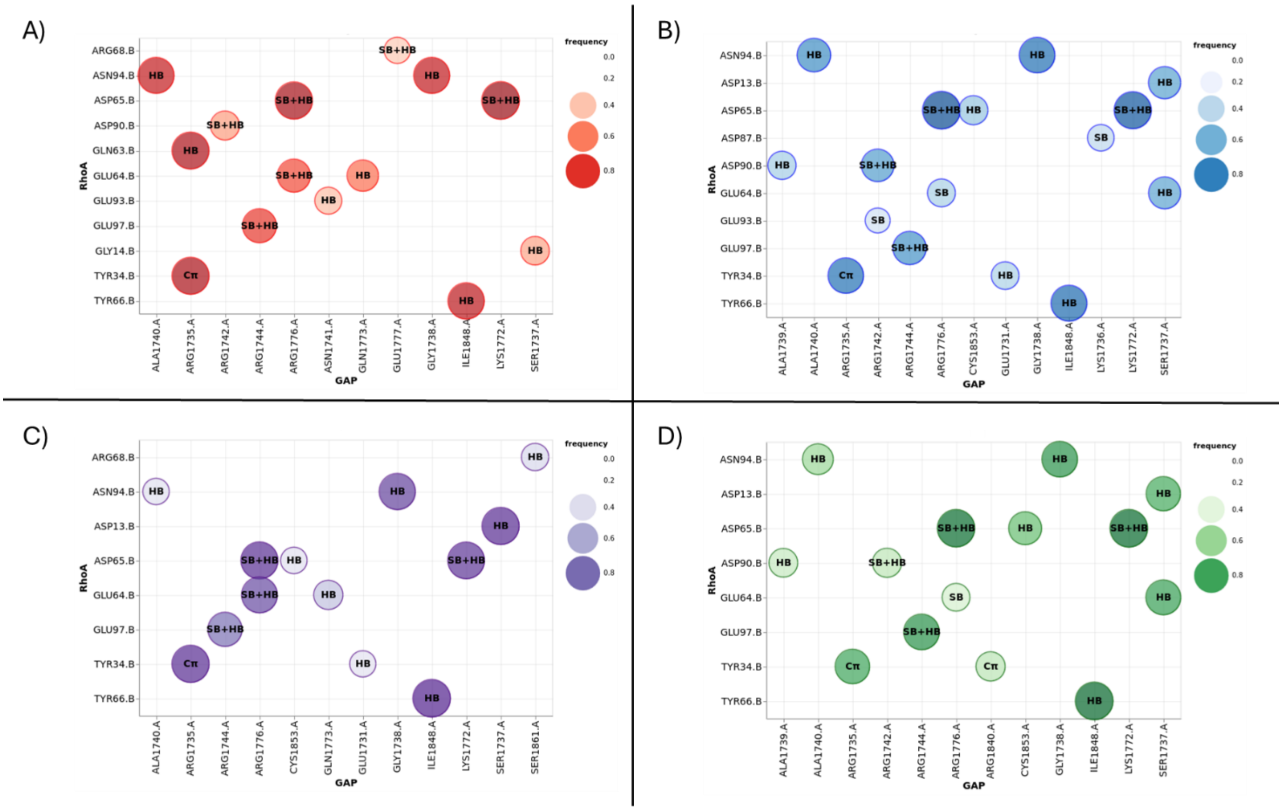

**Figure S10.** Non-covalent interactions (NCIs) established between RhoGAP and RhoA during three 500 ns-long replicas of classical MD simulations. Systems are shown as (a) REA (red), (b) INT3 (blue), (c) INT4 (violet), and (d) PROD (green). HB indicates hydrogen bonds, SB indicates salt bridges, and Cπ denotes cation–π interactions. The frequency of each interaction is indicated by size and color intensity of the cycle, as shown in the legend. NCIs were obtained by using ProLIF [1] a Python library for generating interaction fingerprints in molecular complexes. Additional details are provided in Table S2.

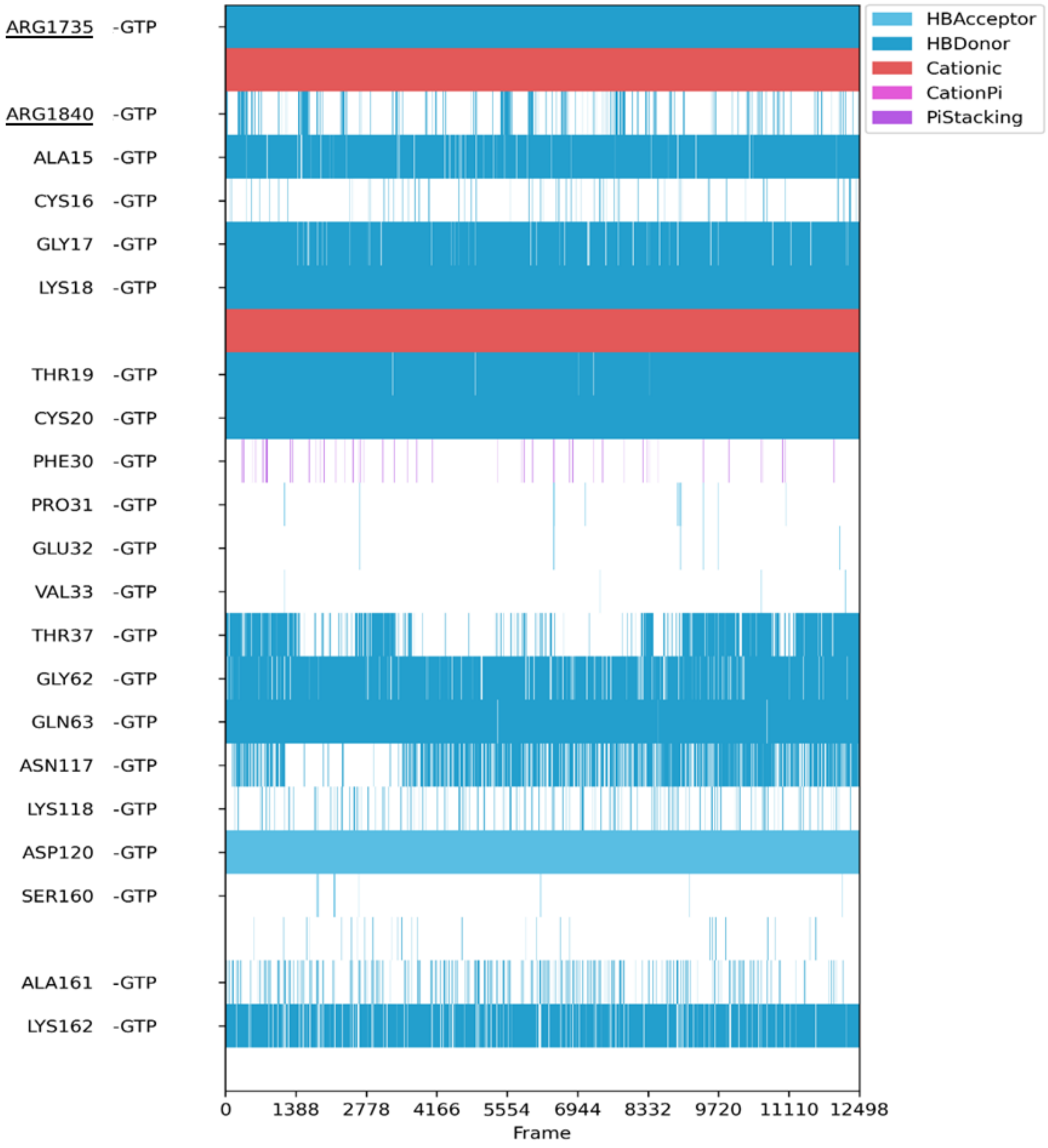


**Figure S11.** Non-covalent interactions (NCIs) analysis between GTP and RhoGAP/RhoA proteins. The graph was generated analysing three 500 ns-replicas of MD simulations with ProLIF python package[1]. Residues belonging to RhoGAP are underlined. Additional details about interactions are provided in Table S3.

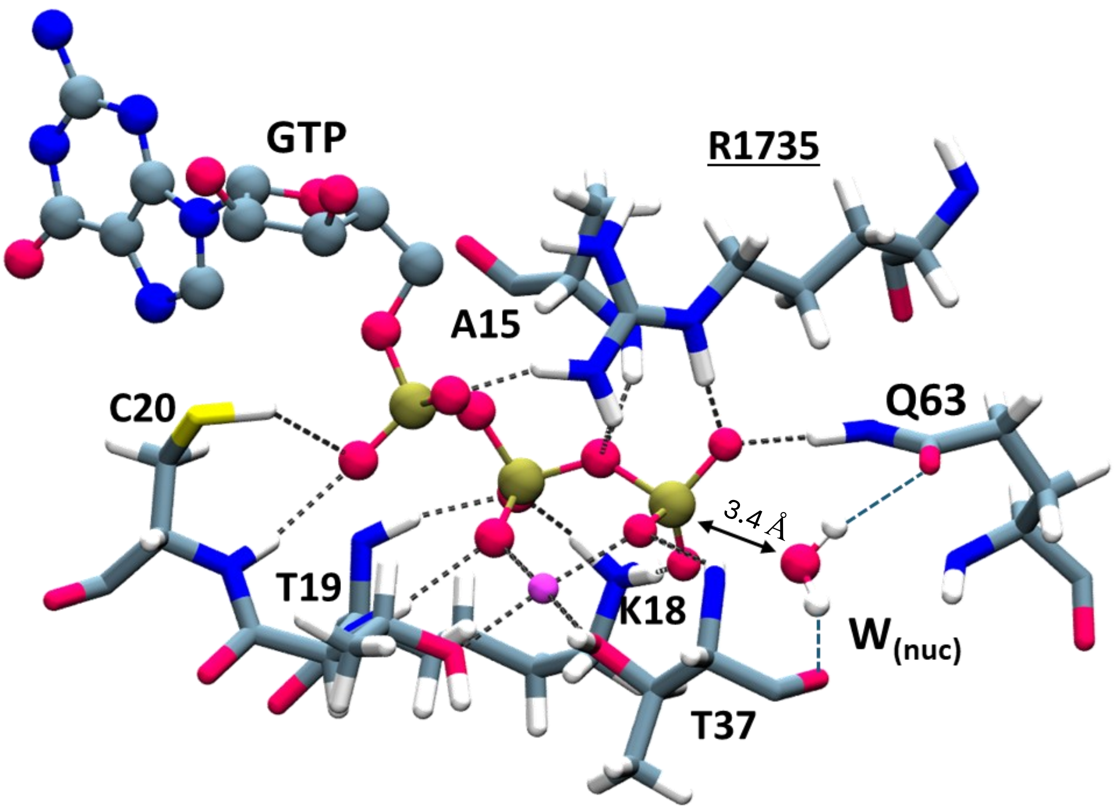


**Figure S12.** Representative frame of classical MD simulation depicting the interactions between the triphosphate moiety of GTP and the surrounding protein residues in the reactant state. Protein residues are shown in sticks, while the GTP is depicted in ball and sticks. Carbon, oxygen, nitrogen, phosphorous and hydrogen atoms are colored in gray, red, blue, brown, and white respectively. $Mg^{2+}$ ion is shown as a magenta van der Waals sphere.

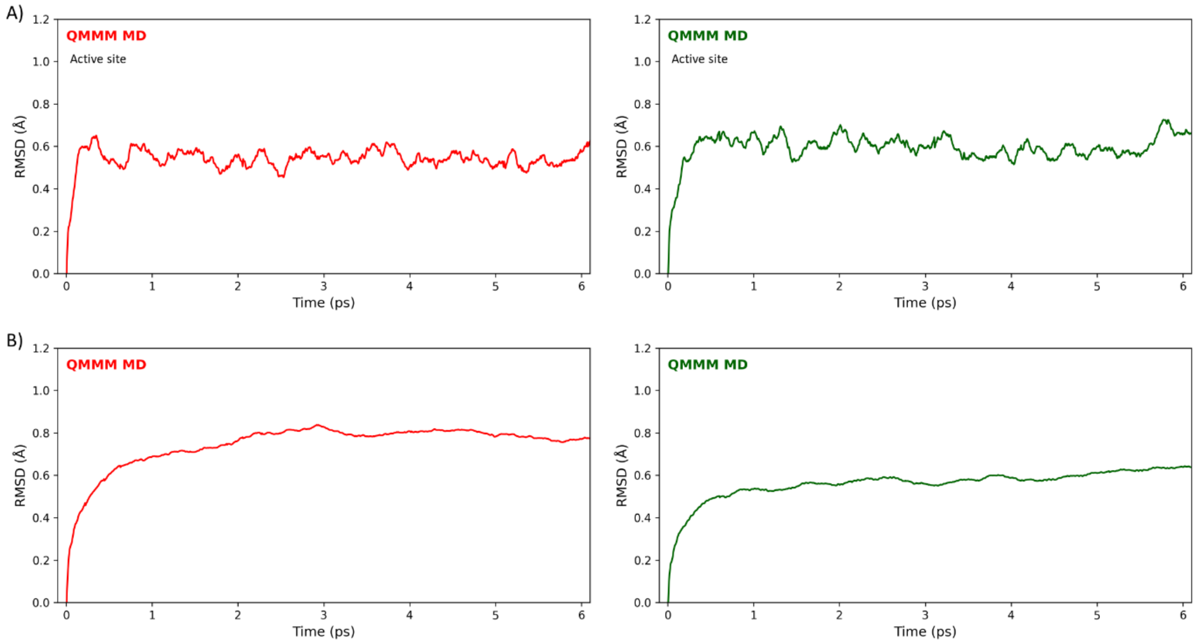


**Figure S13.** Root Mean Square Deviation (RMSD, Å) of the QM region (A) and of the entire system (B) computed over 6 ps-long of QM/MM MD simulations for the REA (red) and PROD (green) systems.

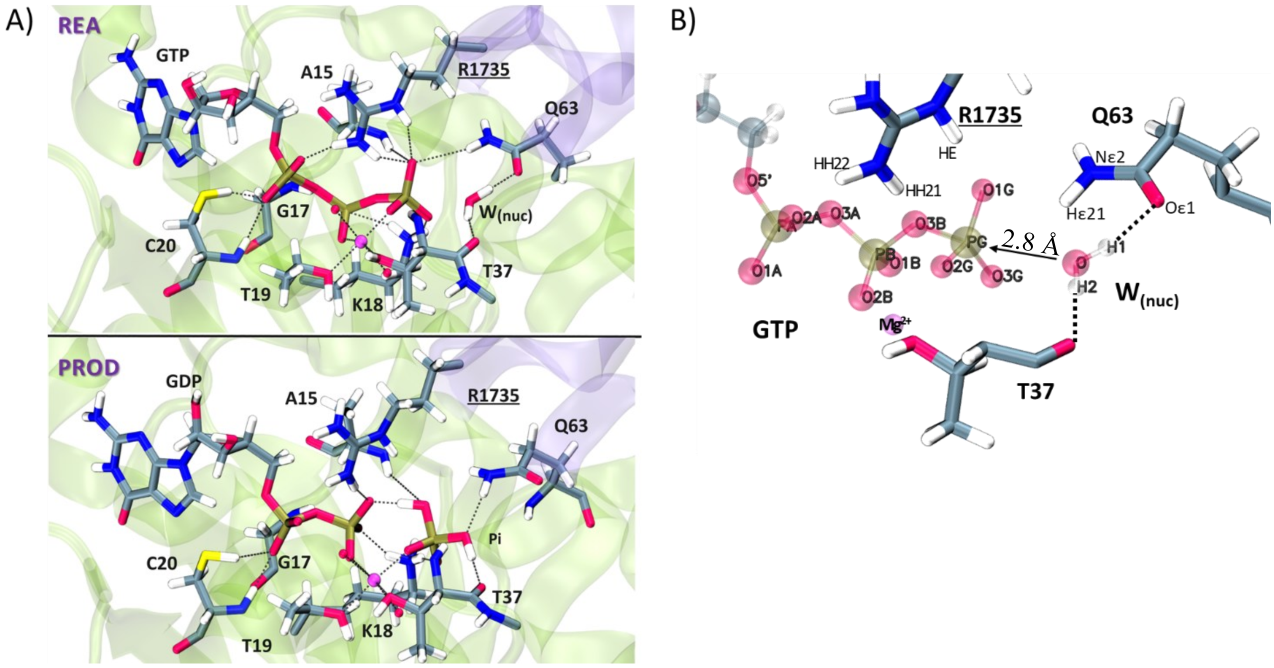


**Figure S14.** A) Most representative frames of QM/MM MD simulation depicting the interactions between the GTP triphosphate moiety and the surrounding protein residues in the reactant (REA) state (top), and in the product (PROD) state (bottom). Hydrogen bonds and metal coordination interactions are indicated with dashed lines. Protein residues and the GTP are shown in sticks. Carbon, oxygen, nitrogen, phosphorous and hydrogen atoms are colored in gray, red, blue, brown, and white, respectively. $Mg^{2+}$ ion is shown as a magenta van der Waals sphere. B) Orientation of the GTP and the nucleophilic water (ball-and-stick) captured in representative frame of QM/MM MD simulations of the reactant state (REA). Further details on the distances are reported in Table S4.

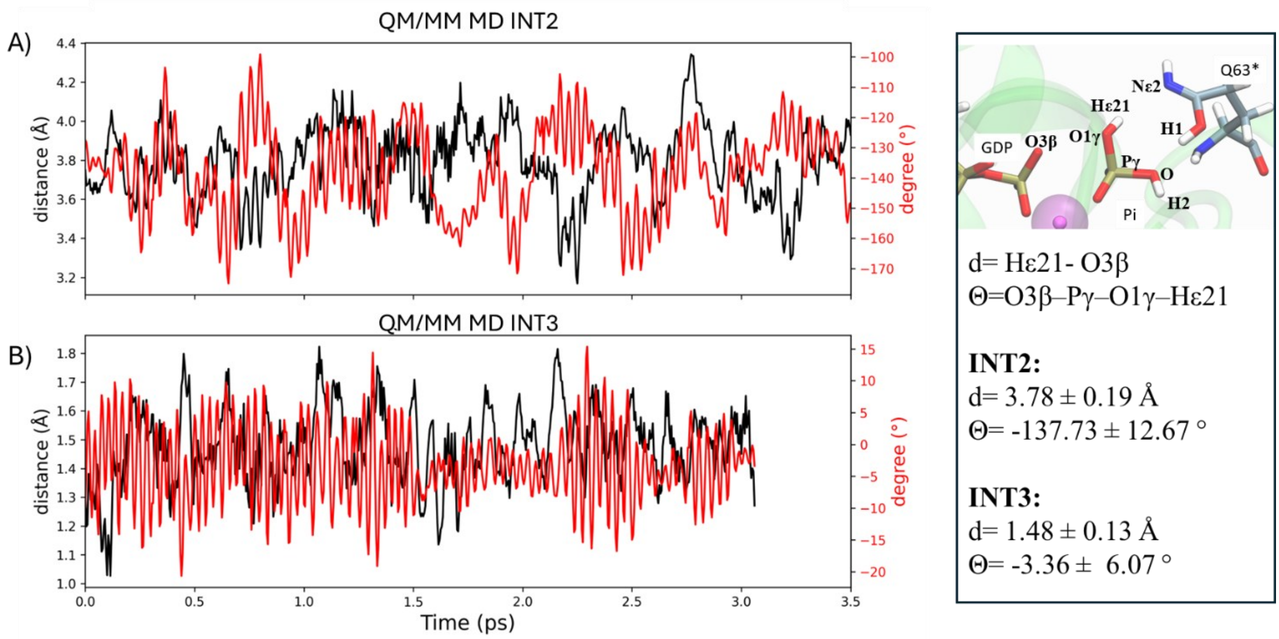


**Figure S15.** Evolution of the dihedral angle defined by O3β:GDP–Pγ:Pi–O1γ:Pi–Hε21:Gln63 (θ, °) and the distance between Hε21 and O3β (d, Å) during QM/MM MD simulations of INT2 (A) and INT3 (B). The inset shows the atom legend and reports the average values and standard deviations for the dihedral angle and the distance.

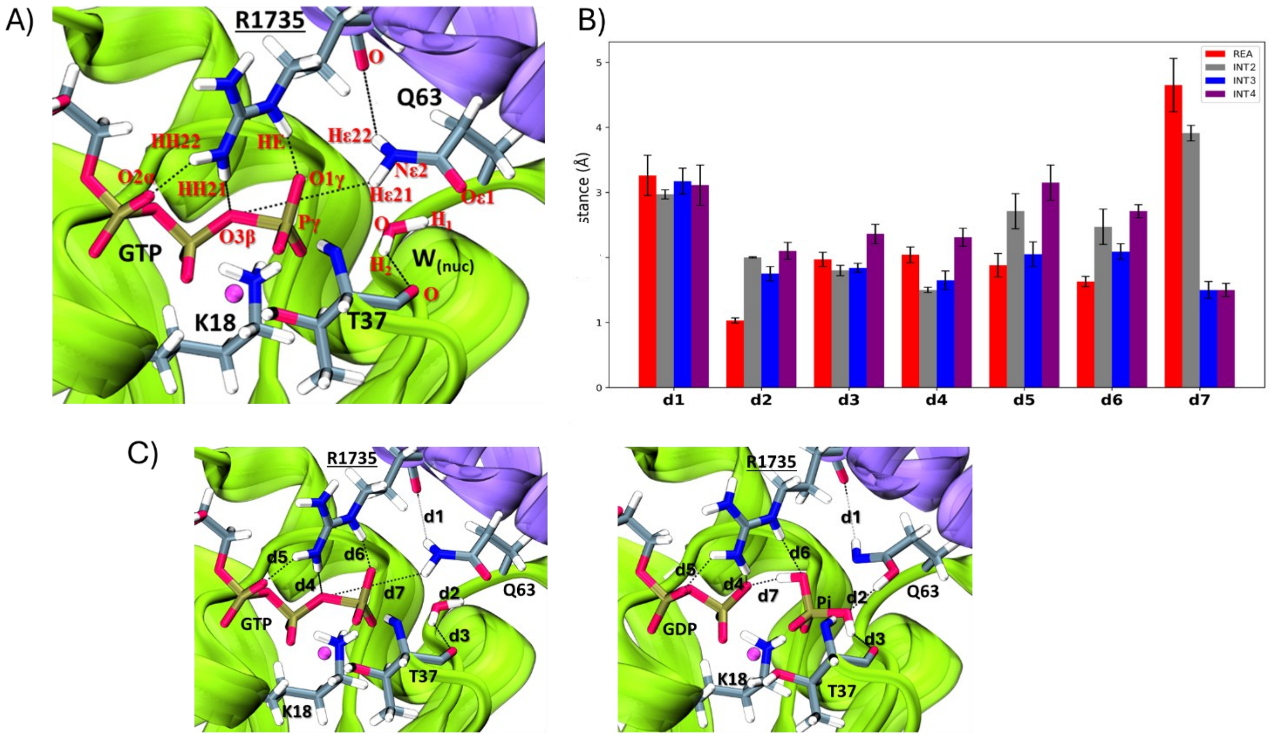


**Figure S16.** A) Active site in the reactant state. Residues are named in black and selected key atoms are highlighted in red. Protein residues and the GTP are shown in sticks. Carbon, oxygen, nitrogen, phosphorous and hydrogen atoms are colored in gray, red and blue, brown, white respectively. $Mg^{2+}$ ion is shown as a magenta van der Waals sphere. B) Set of interatomic distances (Å) for the REA (red), INT2 (grey), INT3 (blue), and INT4 (violet) states, calculated from representative structures of each minimum of reaction coordinate (two from replica 1 and one from replica 2). Error bars report the standard deviation. The distances shown in panel B are defined as follows: d1 = O:Arg1735–Hε22:Gln63, d2 = O:Wnuc–H1:Wnuc, d3 = O:Thr37–H2:Wnuc, d4 = HH21:Arg1735–O3β:GTP, d5 = HH22:Arg1735–O2α:GTP, d6 = HE:Arg1735–O1γ:GTP, d7 = Hε21:Gln63–O3β:GTP. C) Selected interatomic distances in the REA (left) and INT3 (right) states.

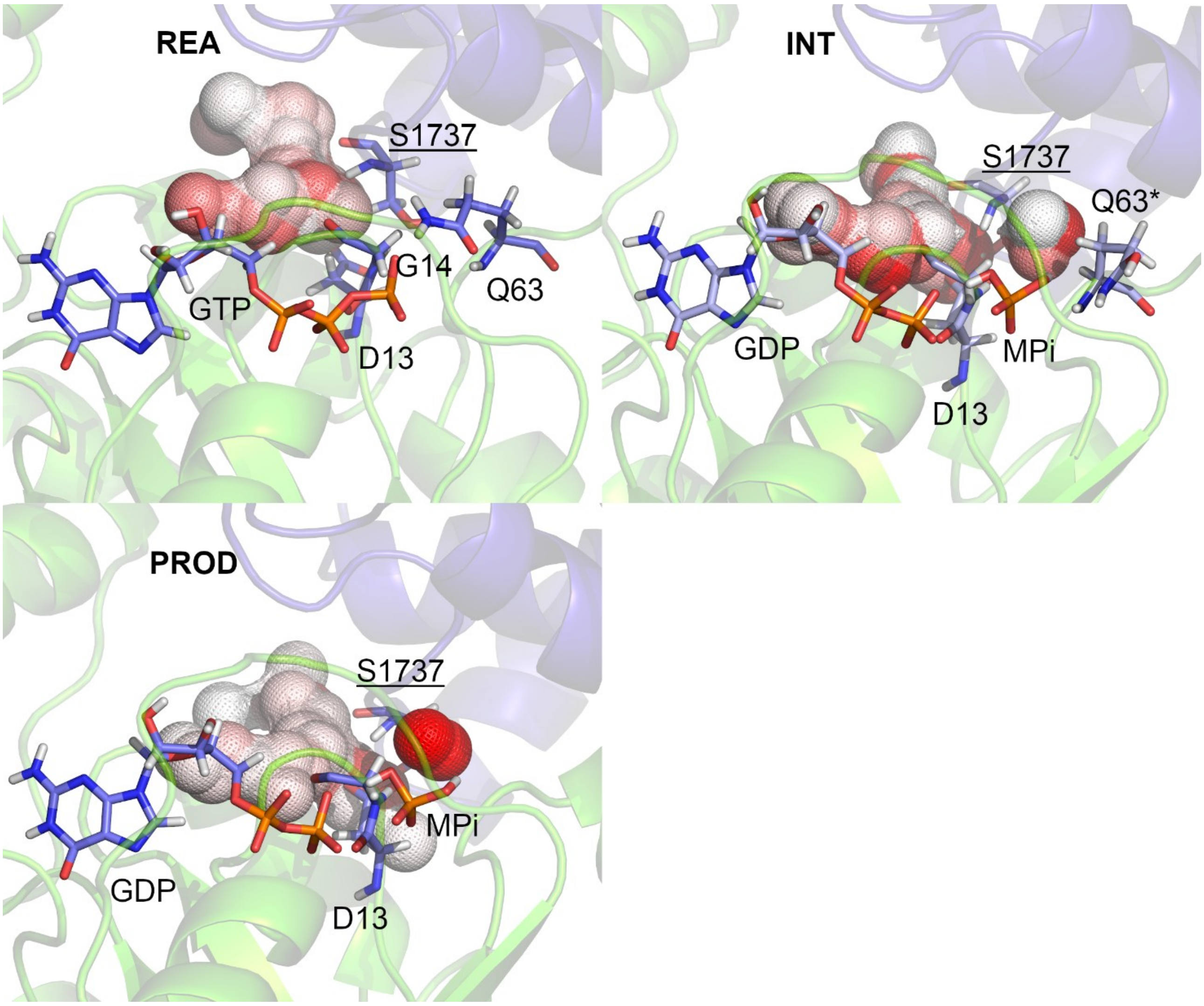


**Figure S17.** Hydration hotspot analysis for REA, INT4, and PROD states. Coloured spheres denote hydration hotspots, defined as regions where water molecules exhibit residence times longer than average. Colour intensity (darker red) reflects an increasing residence time of water. The protein is shown as a transparent cartoon, with Asp13, Gly14, Gln63:RhoA, Ser1737:RhoGAP, GTP, GDP, and MPi depicted as sticks. Carbon, oxygen, nitrogen, phosphorous and hydrogen atoms are colored in violet, red, blue, brown, and white, respectively. $Mg^{2+}$ ion is shown as a magenta van der Waals sphere. Water molecule tracking was performed using AQUA-DUCT v.1.0. [2] on the first 300 ns of the simulations, as detailed in the Methods section of the main text.

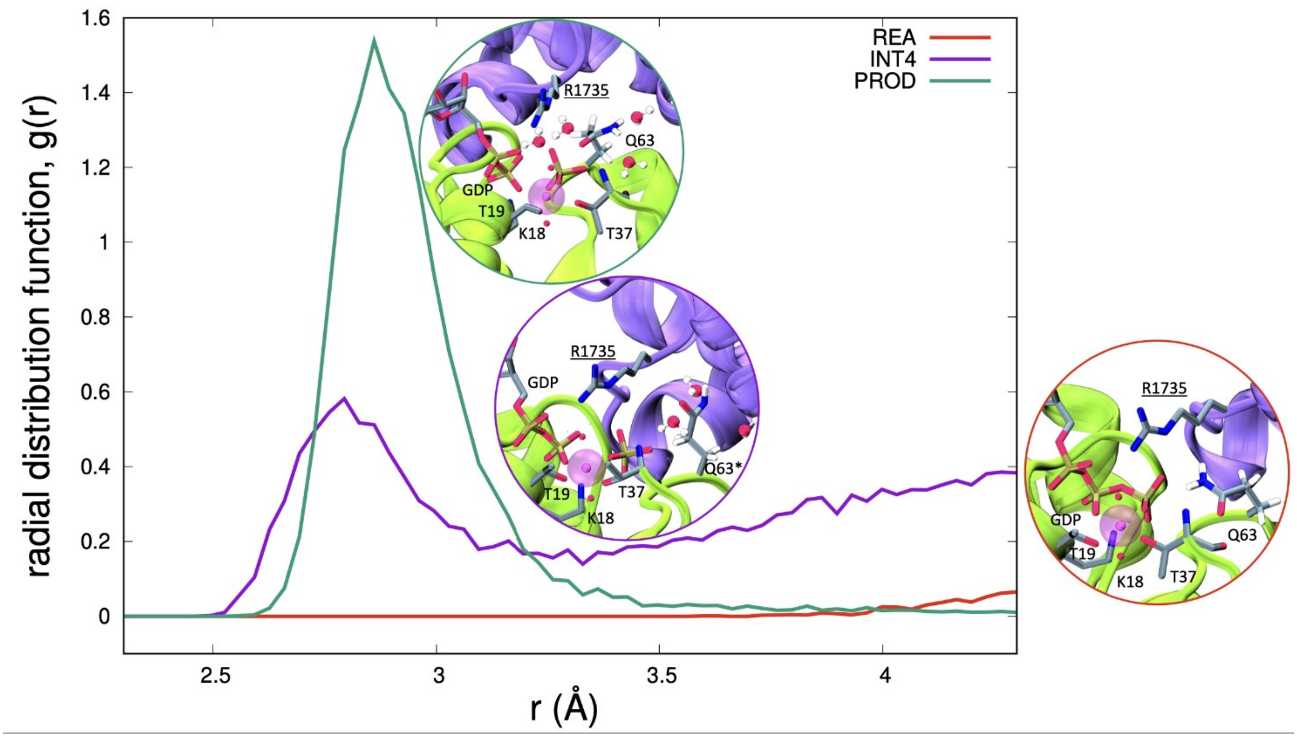


**Figure S18.** Radial distribution function (RDF) computed in REA (red, with red inset), INT4 (purple line, with purple inset) and PROD (green line, with green inset) states. The protein is shown as a cartoon, with relevant species, Gln63/Gln63* and the water molecules in its proximity depicted as sticks. RDF is computed on the first 300 ns of the production run and on a total of 7500 frames. Protein residues and the GTP are shown in sticks. Carbon, oxygen, nitrogen, phosphorous and hydrogen atoms are colored in gray, red, blue, brown, and white, respectively. $Mg^{2+}$ ion is shown as a magenta van der Waals sphere.

*Consistent with water-tracking analyses (Figure S17), the radial distribution function (RDF) of solvent molecules around Gln63 reveals a distinct state-dependent hydration pattern. In the REA state, negligible water density is observed near Gln63, indicating a tightly closed catalytic site. A pronounced RDF peak appears in INT4 and further increases in PROD, reflecting progressive solvent accessibility to the active site. This hydration pattern aligns with the weakening of the RhoGAP:RhoA interface upon reaction completion and confirms that complex dissociation is facilitated after GDP + Pi formation, when water molecules gain access to and destabilize the RhoA:RhoGAP interface.*

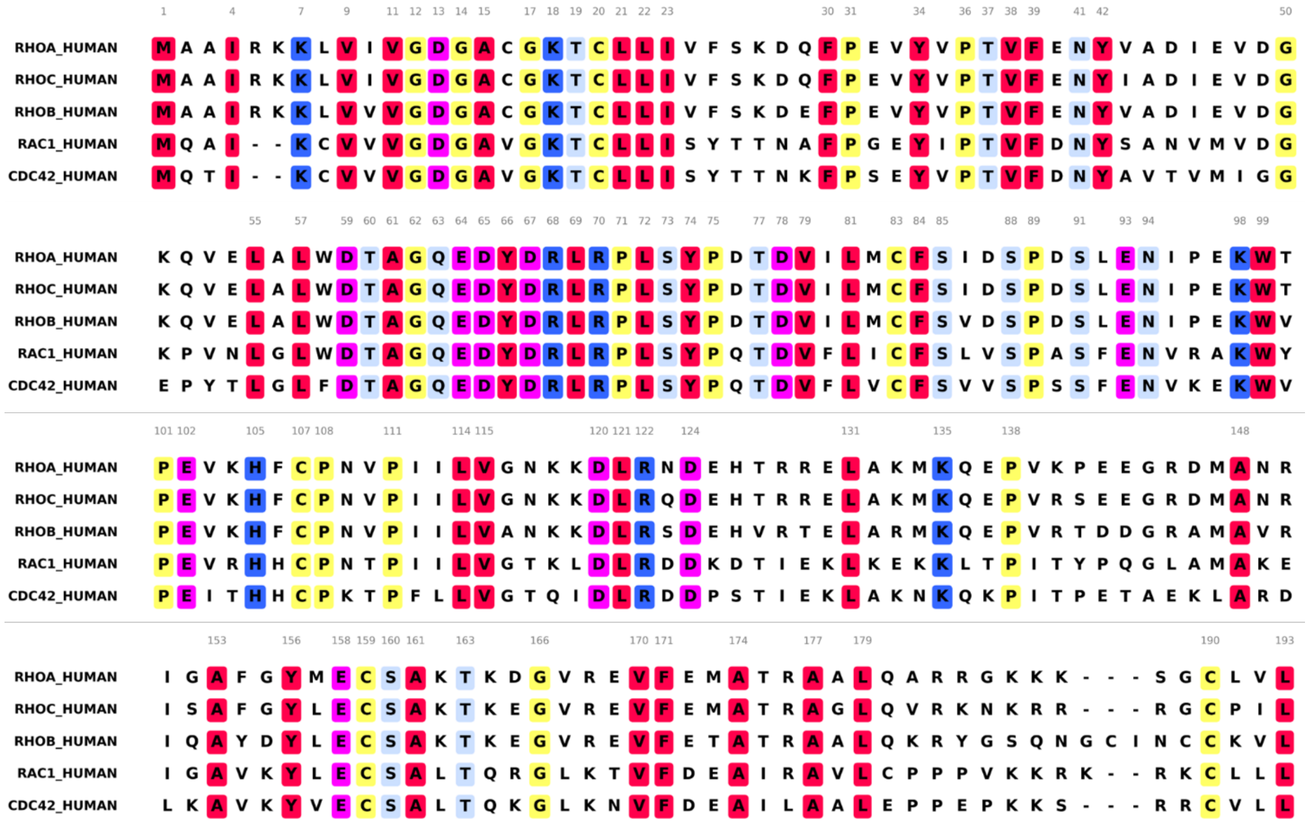

**Figure S19.** Sequence alignment of human Rho family. The colour code, used to highlight conserved residues across all subfamilies is as follows: amino acids with hydrophobic side chains are shown in red, polar uncharged side chains in light blue, positively and negative charged side chains in blue, and magenta, respectively. All other residues in yellow.

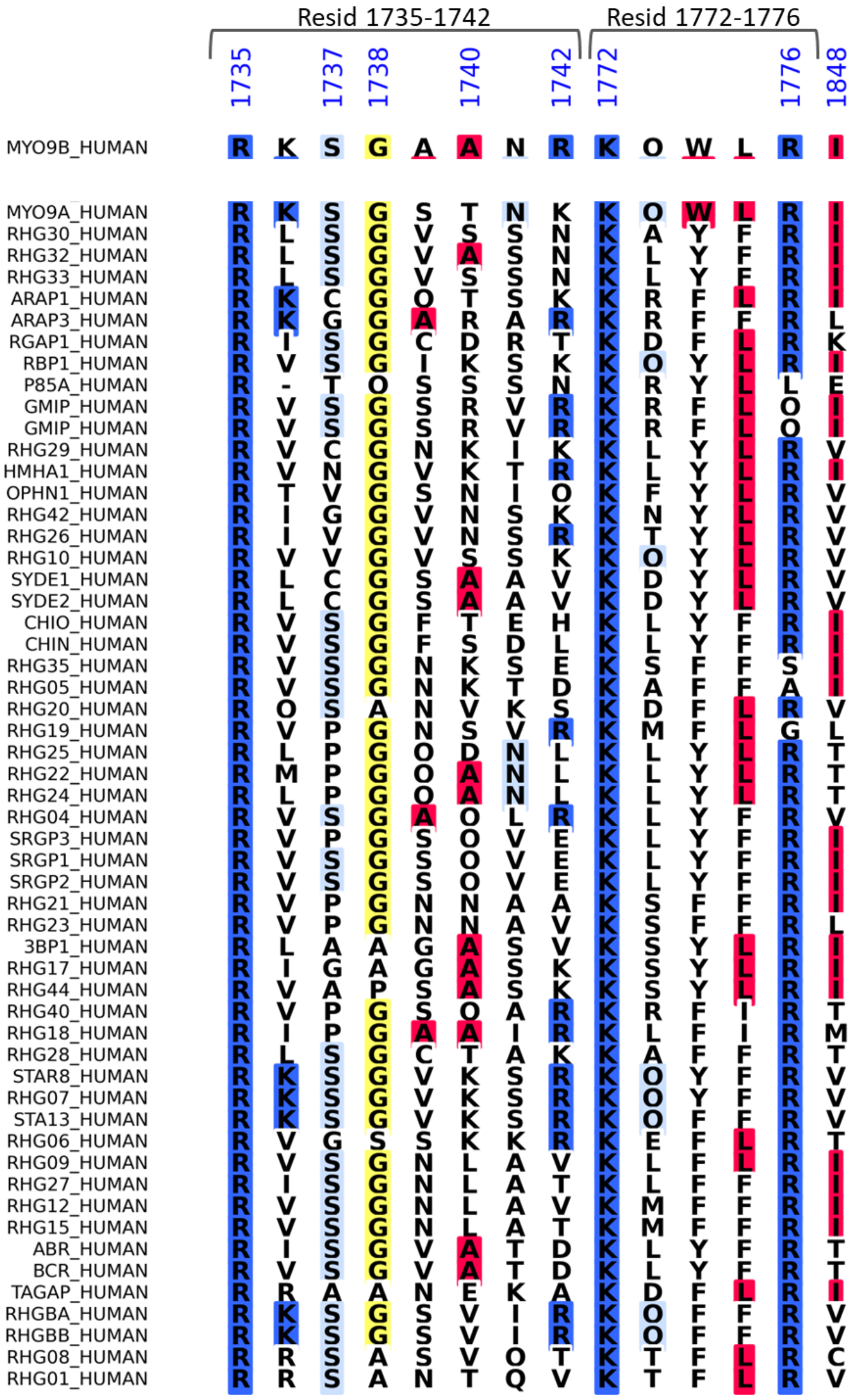

**Figure S20.** Sequence alignment of human GAP residues establishing interactions RhoA protein. Colour codes are the same of Figure S19. At position 1737 Ser is mostly present (54,4%), followed by Pro (16,1%), Cys (7.1%), Gly (7.1%), Ala (5.3%), Val (5.3%), Thr (1,8%) and Asn (1,8%).

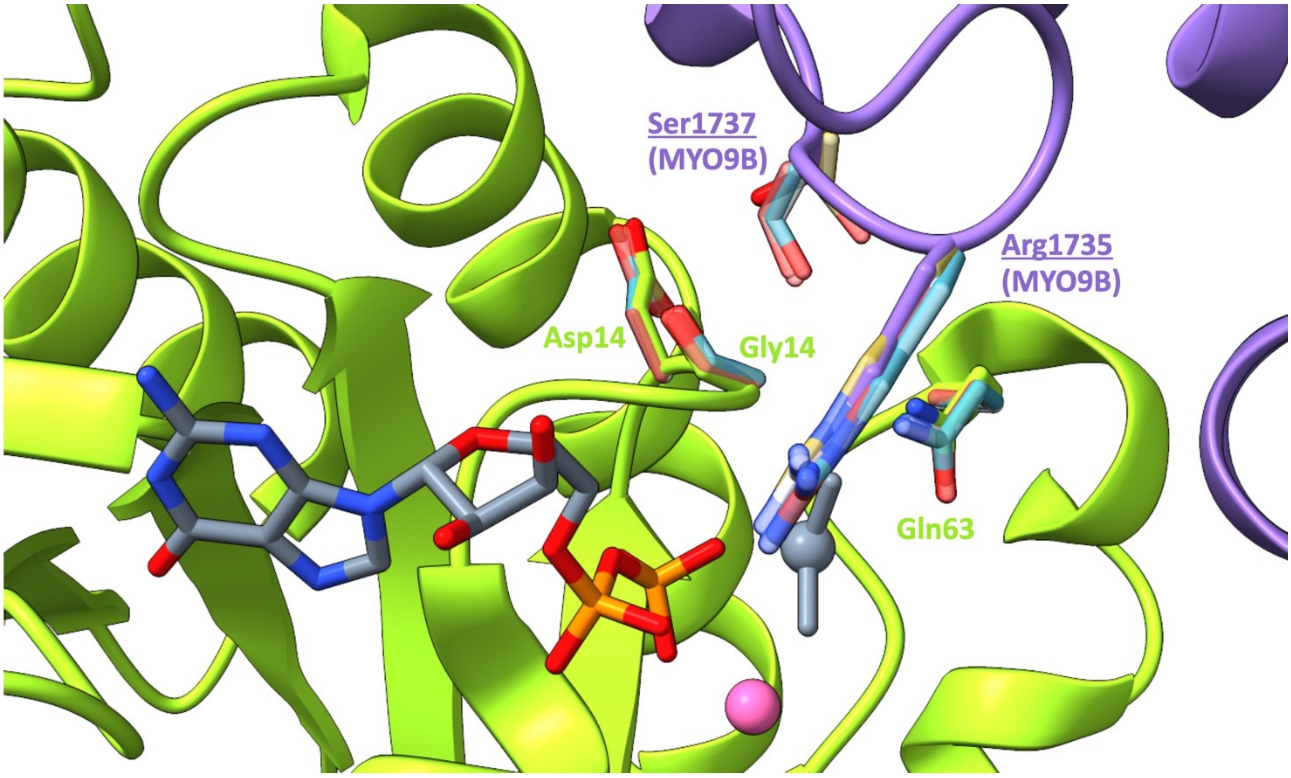


**Figure S21.** Structure of RhoGAP (MYO9B) as deposited in PDB 5HPY. RhoA residues and GAP (MYO9B) are depicted in purple and green, respectively. The GDP and the co-cristallisation element $MgF_3^-$ (in dark gray), are represented as sticks. The superposition of the interacting couples in GAP:RhoA complex Arg1735-Gln63 and Ser1737 are reported for 5 GAP families, namely RGH30, CHIO, SRG1 and ABR whose carbon atoms are colored in yellow, cyan, red and grey, respectively (The not-visible residues are superimposing with other systems). The structures for the GAP:RhoA complex involving GAP proteins RGH30, CHIO, SRG1 and ABR were obtained via AlphaFold3[3] webserver.

**Table S1.** Detailed description of the collective variables (CVs) and parameters used in metadynamics simulations.

| | Simulations time (ps) | CV | Coordinate | Width | MD Time Step (fs) | Time Between Hills (fs) |
|---|---|---|---|---|---|---|
| **MTD1** | 14 | 1 | [O3β:GTP–Pγ:GTP]–[O:Wnuc–Pγ:GTP] | 0.5 a.u. / 0.26 Å | 0.5 | 25 |
| | | 2 | [Hε21–Nε2:Gln63]–[Oε1:Gln63–H1:Wnuc] | 0.5 a.u. / 0.26 Å | 0.5 | 25 |
| **MTD1'** | 8 | 1 | [O3β:GTP–Pγ:GTP]–[O:Wnuc–Pγ:GTP] | 0.5 a.u. / 0.26 Å | 0.5 | 25 |
| | | 2 | [Hε21–Nε2:Gln63]–[Oε1:Gln63–H1:Wnuc] | 0.5 a.u. / 0.26 Å | 0.5 | 25 |
| **MTD2** | 4.5 | 3 | [H1:Gln63*–Nε2:Gln63*] | 0.5 a.u. / 0.26 Å | 0.5 | 25 |
| **MTD3** | 6 | 4 | [H2:Pi–Nε2:Gln63*] | 0.5 a.u. / 0.26 Å | 0.5 | 25 |
| | | 5 | [H1:Gln63*–Oε1:Gln63*]–[H:Gln63*–O:Pi] | 1.0 a.u. / 0.50 Å | 0.5 | 25 |
| **MTD3'** | 7 | 4 | [H2:Pi–Nε2:Gln63*] | 1.0 a.u. / 0.50 Å | 0.5 | 25 |
| | | 5 | [H1:Gln63*–Oε1:Gln63*]–[H:Gln63*–O:Pi] | 1.0 a.u. / 0.50 Å | 0.5 | 25 |
| **MTD4** | 3 | 6 | [H1:Gln63*–O:W3] | 0.5 a.u. / 0.26 Å | 0.5 | 25 |
| | | 7 | [H1:W3–H1: Nε2:Gln63*] | | | |
| **MTD4'** | 5 | 6 | [H1:Gln63*–O:W3] | 0.5 a.u. / 0.26 Å | 0.5 | 25 |
| | | 7 | [H1:W3–H1: Nε2:Gln63*] | | | |
| **MTD4''** | 4 | 6 | H1:Gln63*–O:W3] | 0.5 a.u. / 0.26 Å | 0.5 | 25 |
| | | 7 | [H1:W3–H1: Nε2:Gln63*] | | | |

**Table S2.** Interaction frequencies between RhoGAP and RhoA, grouped by residue, calculated from classical MD simulations of the REA (red), INT3 (blue), INT4 (violet), and PROD (green) systems. For each interaction, the persistence over the 500 ns MD simulation normalized to 1 (Freq), is reported. The arginine finger is shown in bold. Interactions with a frequency lower than 0.45 are not shown.

| RhoGAP | RhoA | REA | INT3 | INT4 | PROD |
|---|---|---|---|---|---|
| Residue | | Freq | | | |
| LYS1772 | ASP65 | 1.0 | 1.0 | 0.9 | 1.0 |
| ARG1776 | ASP65 | 1.0 | 1.0 | 1.0 | 1.0 |
| ARG1735 | TYR34 | 1.0 | 0.9 | 1.0 | 0.8 |
| ARG1735 | GLN63 | 1.0 | - | - | - |
| ILE1848 | TYR66 | 0.9 | 0.9 | 1.0 | 1.0 |
| GLY1738 | ASN94 | 0.9 | 0.9 | 0.9 | 0.9 |
| ALA1740 | ASN94 | 0.9 | 0.8 | 0.4 | 0.6 |
| ARG1744 | GLU97 | 0.8 | 0.7 | 0.8 | 0.9 |
| ARG1776 | GLU64 | 0.7 | 0.5 | 0.9 | 0.4 |
| GLN1773 | GLU64 | 0.6 | - | 0.5 | - |
| ARG1742 | ASP90 | 0.5 | 0.7 | - | 0.5 |
| SER1737 | GLY14 | 0.5 | - | - | - |
| ASN1741 | GLU93 | 0.5 | - | - | - |
| GLU1777 | ARG68 | 0.5 | - | - | - |
| SER1737 | ASP13 | - | 0.7 | 1.0 | 0.8 |
| SER1737 | GLU64 | - | 0.7 | - | 0.8 |
| CYS1853 | ASP65 | - | 0.5 | - | 0.7 |
| ALA1739 | ASP90 | - | 0.5 | - | 0.5 |
| ARG1840 | TYR34 | - | - | - | 0.5 |

**Table S3.** Non covalent interactions (NCIs) between GTP and RhoGAP/RhoA proteins in the reactant state. The persistence along the 500 ns MD simulation trajectory normalized to 1 (Freq) is reported. Residues belonging to RhoGAP are underlined. The hydrogen-bond labelled (d) indicates that the donor atom of the hydrogen belongs to GTP.

| **Residue** | **Ligand** | **Interaction type** | **Freq** |
|---|---|---|---|
| <u>ARG1735</u> | GTP | H-bond | 1.0 |
| <u>ARG1735</u> | GTP | Salt bridge | 1.0 |
| LYS18 | GTP | H-bond | 1.0 |
| LYS18 | GTP | Salt bridge | 1.0 |
| CYS20 | GTP | H-bond | 1.0 |
| ASP120 | GTP | H-bond (d) | 1.0 |
| GLN63 | GTP | H-bond | 1.0 |
| THR19 | GTP | H-bond | 1.0 |
| ALA15 | GTP | H-bond | 1.0 |
| GLY17 | GTP | H-bond | 1.0 |
| GLY62 | GTP | H-bond | 0.9 |
| LYS162 | GTP | H-bond | 0.9 |
| ASN117 | GTP | H-bond | 0.6 |
| THR37 | GTP | H-bond | 0.5 |
| <u>ARG1840</u> | GTP | H-bond | 0.4 |

**Table S4.** Distances d (Å) between RhoGAP and RhoA residues and the GDP in REA and GDP + Pi in PROD, respectively. The HE21 atom, which belongs to Gln63 in the reactants and to Pi in the products, is marked in bold. All distances (d) were calculated over 6 ps-long QM/MM MD simulation trajectories and are reported along with their standard deviation (sd). Distances larger than 2.45 Å are highlighted in red.

| RhoGAP:Rhoa | | Substrate | REA | | PROD | |
|---|---|---|---|---|---|---|
| Resid | atom | atom | <d> | sd | <d> | sd |
| MG | $Mg^{2+}$ | O2G | 2.06 | 0.08 | 2.05 | 0.08 |
| MG | $Mg^{2+}$ | O2B | 2.04 | 0.08 | 2.03 | 0.07 |
| ARG1735 | HE | O1G | 1.67 | 0.12 | 2.58 | 0.26 |
| ARG1735 | HH22 | O2A | 1.97 | 0.25 | 2.53 | 0.23 |
| ARG1735 | HH21 | O3B | 2.26 | 0.36 | 1.67 | 0.17 |
| LYS18 | HZ2 | O3G | 1.61 | 0.12 | 1.71 | 0.14 |
| LYS18 | HZ1 | O1B | 1.77 | 0.16 | 1.66 | 0.13 |
| THR19 | H | O2B | 2.29 | 0.13 | 2.37 | 0.15 |
| CYS20 | H | O1A | 2.20 | 0.15 | 2.04 | 0.12 |
| **GLN63(Pi)** | **HE21** | **O1G** | **2.02** | **0.20** | **1.12** | **0.14** |
| LYS18 | H | O1B | 2.29 | 0.22 | 2.45 | 0.20 |
| THR37 | H | O2G | 2.34 | 0.21 | 2.06 | 0.18 |
| GLY17 | H | O1B | 2.40 | 0.27 | 2.63 | 0.32 |
| ALA15 | H | O3B | 2.20 | 0.14 | 2.10 | 0.12 |
| ALA15 | HA | O3A | 2.54 | 0.17 | 2.72 | 0.20 |
| GLY62 | H | O3G | 2.37 | 0.22 | 2.25 | 0.23 |
| GLY17 | HA2 | H8 | 2.45 | 0.21 | 2.52 | 0.26 |
| GLY17 | H | O3A | 2.55 | 0.27 | 2.32 | 0.19 |
| CYS20 | HG | O1A | 2.20 | 0.28 | 3.43 | 1.03 |
| LYS18 | HG3 | O1B | 2.61 | 0.22 | 2.62 | 0.24 |
| LYS162 | H | O6 | 2.57 | 0.26 | 2.33 | 0.13 |
| THR19 | HB | O1A | 2.59 | 0.22 | 2.58 | 0.18 |
